# Quantifying the susceptibility of polygenic scores to ancestry stratification

**DOI:** 10.64898/2025.12.04.692430

**Authors:** Jennifer Blanc, Walid Mawass, Jeremy J. Berg

## Abstract

Polygenic scores aim to predict phenotypes from genetic data, yet they remain vulnerable to spurious correlations arising from environmental variation that covaries with population structure. While standard methods like Principal Component Analysis (PCA) and Linear Mixed Models (LMMs) mitigate this, quantifying the residual risk for specific applications remains challenging. Here, we develop a theoretical framework that quantifies the proportion of genetic variance in a GWAS panel explained by an external ancestry gradient (*H*), providing a direct measure of stratification susceptibility. We show that this baseline risk is amplified by the ascertainment process itself, which creates a directional bias (Φ) that is particularly strong for variants with intermediate probabilities of ascertainment. Applying this framework to the UK Biobank, we find that while uncorrected susceptibility is drastically higher in diverse cohorts, PCA correction effectively flattens this disparity. We observe that the residual susceptibility (*H*^′^) in corrected diverse panels is often comparable to, or marginally lower than, that found in restricted homogeneous subsets, suggesting that sample diversity need not compromise stratification control. However, for both study designs, residual structure often remains just above or indistinguishable from the theoretical limit of detection. Because even undetectable levels of structure can accumulate to produce significant bias in highly polygenic scores, we introduce a diagnostic to calculate the critical magnitude of environmental confounding required to explain an observed signal. Using this diagnostic, we find that both the well-known divergence in height scores between Sardinia and mainland Europe and novel signals of divergence in systolic blood pressure scores within the British Isles appear relatively robust to residual stratification, albeit for different reasons. While the Sardinia signal would require moderate-to-strong environmental confounding to align with a vanishingly small residual ancestry axis, the systolic blood pressure signals would require implausibly large environmental effects to be explained as artifacts.

## 1 Introduction

Polygenic scores (PGS) aim to aggregate the additive effects of genetic variants, serving as a predictor of genetic liability for complex traits. Ideally, this allows for phenotypic prediction based solely on an individual’s genotype. However, the effect size estimates underlying these scores are derived from population-based genome-wide association studies (GWAS), where genetic variation and environmental factors often covary with population structure. If left uncorrected, these correlations result in systematically biased effect size estimates. When these biases are aggregated across thousands of variants in a prediction panel, they can recapitulate the original patterns of population structure, confounding inferences about evolutionary history [1, 2, 3] and limiting the transferability of PGS predictions [4].

Confounding due to population structure has been a persistent concern since the introduction of polygenic scores [5]. To mitigate this, the field has converged on two statistical correction strategies: Principal Component Analysis (PCA), which models and removes axes of variation as fixed effects [6], and Linear Mixed Models (LMMs), which model the genetic background as a random effect [7, 8]. Biobank-scale analyses often employ a hybrid approach, including top principal components (PCs) as fixed effects alongside an LMM [9, 10]. This represents a pragmatic heuristic: explicitly remove the strongest axes of variation via PCA and rely on the LMM to dampen residual confounding. However, the success of this strategy is contingent on the quality of the fixed effects. It requires not only that the top genetic PCs are accurately estimated from the data [11, 3], but also that the relevant environmental confounder falls within the subspace spanned by these estimated axes. If these conditions are not met, the confounder falls into the residual genetic background, where protection depends on LMM assumptions about the correct amount of shrinkage along different eigenvectors [12, 13, 14]. This parametric uncertainty means that even PGS constructed from “corrected” effect size estimates may retain bias along some axes [3, 15].

Given the parametric uncertainties of statistical correction, researchers also employ bias avoidance strategies. Family-based association studies represent the theoretical gold standard in this regard, as they eliminate stratification by relying solely on within-family variation, which is orthogonal to population structure [16]. Empirical comparisons confirm that population-based estimates are often inflated, either by indirect effects or by stratification, relative to their within-sibship counterparts [17, 18, 19, 20]. However, the scarcity of large family-based cohorts necessitates the continued use of unrelated population samples for many analyses. To avoid bias in population samples, researchers often aim to ensure that genetic variation within the GWAS panel is independent of the ancestry gradients found in the prediction set. In practice, this is attempted either by restricting analysis to ostensibly homogeneous samples (e.g., ancestry subsets in the UK Biobank [21]) or by using GWAS panels that are evolutionarily diverged from the target prediction panel [5, 22, 3]. However, these strategies are not panaceas: reduced sample size and evolutionary divergence come at the cost of statistical power [23], and because no human population is truly panmictic, stratification concerns persist at every scale [24]. Furthermore, the structural independence required by these designs is often assumed based on broad features of demographic history or geographic separation, rather than explicitly quantified.

Finally, LD score regression (LDSC) has become a standard diagnostic tool for identifying confounding in GWAS summary statistics [25]. While an intercept greater than one signals residual stratification, it does not offer a remedy for PGS construction. The intercept can be used to rescale summary statistics, but like the original genomic control approach [26], this rescaling mitigates test statistic inflation without removing the systematic ancestry-associated bias from the effect size estimates themselves. The utility of LDSC also depends on the availability of reference LD scores that accurately capture the correlation structure of the GWAS sample. Because patterns of linkage disequilibrium are determined by specific demographic histories, the utility of LDSC is limited to samples for which ancestry-matched reference data are available. Thus, while LDSC represents a valuable post-hoc diagnostic, we currently lack a direct framework to quantify exactly how susceptible a polygenic score remains to stratification along a specific external ancestry gradient.

To move beyond reliance on global diagnostics and independence assumptions, we develop a framework that quantitatively assesses the susceptibility of a polygenic score to confounding along specific, researcher-defined ancestry gradients. Building on our earlier work [3], we first introduce a statistic, *H*, that estimates the proportion of genetic variance in a GWAS panel explained by an externally defined ancestry gradient. We demonstrate that this baseline susceptibility is amplified by the ascertainment process itself, which generates a directional bias that creates a spurious correlation between causal effects and population structure. We then introduce a method to evaluate the adequacy of population structure estimators (such as principal components) by calculating the residual susceptibility, *H*^′^, remaining after correction. Applying this framework to the harmonized Human Genome Diversity Panel–Thousand Genomes dataset (HGDP1KG) and the UK Biobank (UKB) [27, 28], we empirically evaluate the trade-offs between sample diversity and homogeneity. We find that while uncorrected susceptibility (*H*) is often higher in diverse panels, robust PC correction effectively neutralizes this disparity. Consequently, the residual susceptibility (*H*^′^) in the full UKB cohort is comparable to—and in some cases marginally lower than—that achieved in the ostensibly homogeneous “White British” subset, though in practice the absolute differences in protection between these strategies are small. Finally, we perform polygenic score association tests for 17 phenotypes to demonstrate the practical utility of our diagnostic. We show how *H*^′^ can be used to bracket the risk of stratification bias by calculating the critical magnitude of environmental confounding required to generate an observed signal. Applying this to empirical data, we show that while the well-known divergence in height scores between Sardinia and mainland Europe remains susceptible to plausible levels of residual confounding, novel signals of divergence in systolic blood pressure within the British Isles appear highly robust.

## 2 Theory

In this section, we develop a formal framework to quantify the susceptibility of polygenic score association tests to stratification bias. Our analysis focuses on the covariance between a polygenic score and a specific, researcher-defined axis of population structure in the prediction panel—the “test vector.” As noted by [3], bias in this statistic arises not from global population structure per se, but from the specific projection of this test vector onto the GWAS panel. To apply this insight, we use a generative framework (fully specified in Supplementary Text section S1.1) that partitions genetic variation into components aligned with and orthogonal to this target axis. We then derive the expected bias in the polygenic score test statistic, accounting for two forms of amplification due to ascertainment bias, and define metrics to evaluate the efficacy of principal component analysis (PCA) in mitigating this risk.

### 2.1 Prediction panel genotypes

First, we consider the structure of the prediction panel genotypes. The analysis begins with the researcher specifying an *N* × 1 “test vector” *t*—such as a continuous geographic coordinate, a group membership indicator, a continuous ancestry estimate, or an estimated sample age [29, 30, 31, 32, 33]—which positions prediction panel individuals along an observed axis of interest. We standardize this vector such that *Var*(*t*) = 1.

We model this observed test vector as the sum of three components:

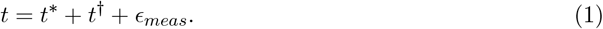

Here, *t*^∗^ represents the component of ancestry variation in *t* that overlaps with that of the GWAS panel due to shared demographic history. The term *t*^†^ represents structured ancestry variation specific to the prediction panel (i.e., orthogonal to the GWAS population structure), and *ϵ*_*meas*_ represents measurement noise independent of ancestry variation.

We then model the mean-centered *N* × *L* prediction panel genotype matrix, **X**, as a function of these latent coordinates:

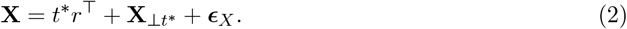

The vector *r* captures the systematic variation in expected genotype associated with the shared structure *t*^∗^. The term *t*^∗^*r*^⊤^ therefore captures the specific component of genetic variation that is both structured by the test vector and shared with the GWAS panel. The residual term **X**_⊥*t*_∗ aggregates all other genetic variation that is structured by the demographic history, including both that associated with *t*^†^ and other axes of variation which may or may not be shared with the GWAS panel, while ***ϵ***_*X*_ represents random sampling noise. We assume that 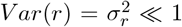, and scale genotypes such that *Var*(*X*_*ℓ*_) = 1.

### 2.2 GWAS panel genotypes

We assume a parallel generative structure for the mean-centered *M* ×*L* GWAS panel genotypes **G**. Specifically, we model the genotypes as arising from the same vector of allelic deviations *r* defined in the prediction panel:

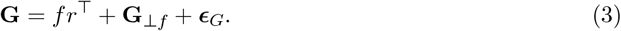

Here, *f* is the unobserved “target axis” representing the position of GWAS individuals along the ancestry gradient defined by the allelic weights *r*. Unlike the test vector *t*, which is an observed variable in the prediction panel, *f* is a latent property of the GWAS sample. Its variance, 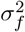, reflects the extent to which the shared ancestry gradient is present in the GWAS panel. Because the test vector *t* is standardized to have unit variance, 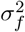 measures the variance in ancestry within the GWAS panel along the target axis relative to the variance along that same axis in the prediction panel. We define *H* as the total variance in the GWAS panel genotypes explained by this target axis:

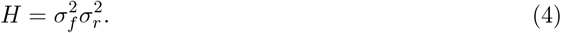

The term **G**_⊥*f*_ captures all other structured variation in the GWAS panel (i.e., other population structure) that is orthogonal to *f* .

### 2.3 Phenotypes

We assume the phenotype vector *y* for the GWAS individuals is generated by:

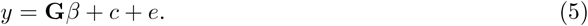

Here, *β* represents the additive effects of the set ℂ of causal variants, which are sampled from an unknown distribution with density *p* (*β*). The vector *c* represents ancestry-associated environmental confounders. Consistent with our genotype decomposition, we partition the confounding variation into a component aligned with the target axis *f* and a component orthogonal to it,

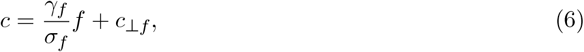

where *γ*_*f*_ represents the magnitude of environmental confounding specifically aligned with the target ancestry axis, and *c*_⊥*f*_ represents confounding aligned with other axes of structure (i.e., **G**_⊥*f*_). Notably, equation (6) represents a purely statistical partition induced by the choice of test vector, *t*, and does not imply that an individual’s ancestry causes their environment.

### 2.4 Polygenic score association test

We test the null hypothesis of no association between the test vector *t* and the prediction panel polygenic effects *z* = **X***β*. We define the target estimand, *q*, as the sample covariance between the test vector and the total polygenic effect:

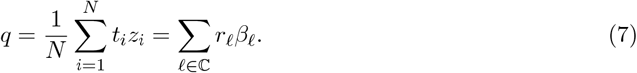

Under the null, 𝔼 [*q*] = 0. We assume that this null holds, and consider the impact of the confounder, *c*, on the test in this case. In practice, variants are ascertained for inclusion based on their estimated effect sizes (e.g., 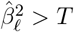, where *T* is determined by a choice of p value significance threshold). Let 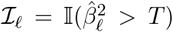) be the indicator variable for whether a variant is ascertained. The estimated covariance can then be written as a sum across all *L* sites measured in the GWAS:

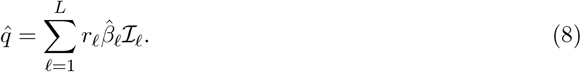

We model the estimated effect sizes as the causal effects plus systematic stratification bias and random noise:

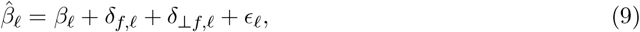

where *δ*_*f,ℓ*_ = *γ*_*f*_ *σ*_*f*_ *r*_*ℓ*_ is the systematic bias along the target axis *f* [3], and *δ*_⊥*f,ℓ*_ is the total bias along background axes orthogonal to *f* (i.e., attributable to **G**_⊥*f*_).

We define the baseline ascertainment probability, *S*(*β*_*ℓ*_) = *P* ( ℐ_*ℓ*_ = 1), as the power to detect a variant with causal effect *β*_*ℓ*_ in the absence of stratification bias, with *S*^′^(*β*_*ℓ*_) and *S*^′′^(*β*_*ℓ*_) giving the first and second derivatives with respect to *β*_*ℓ*_ (Figure 1a-c). Also, let 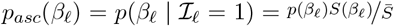, be the density on causal effect sizes conditional on ascertainment, where 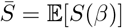 is the average statistical power over all variants. In the Supplementary Text Section S1.2, we derive the expected standardized bias to second order in the ascertainment probability, showing that the bias depends on the fraction of variance explained by the target axis *H*, amplified by two distinct ascertainment effects,

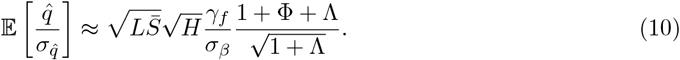

**Figure 1:**
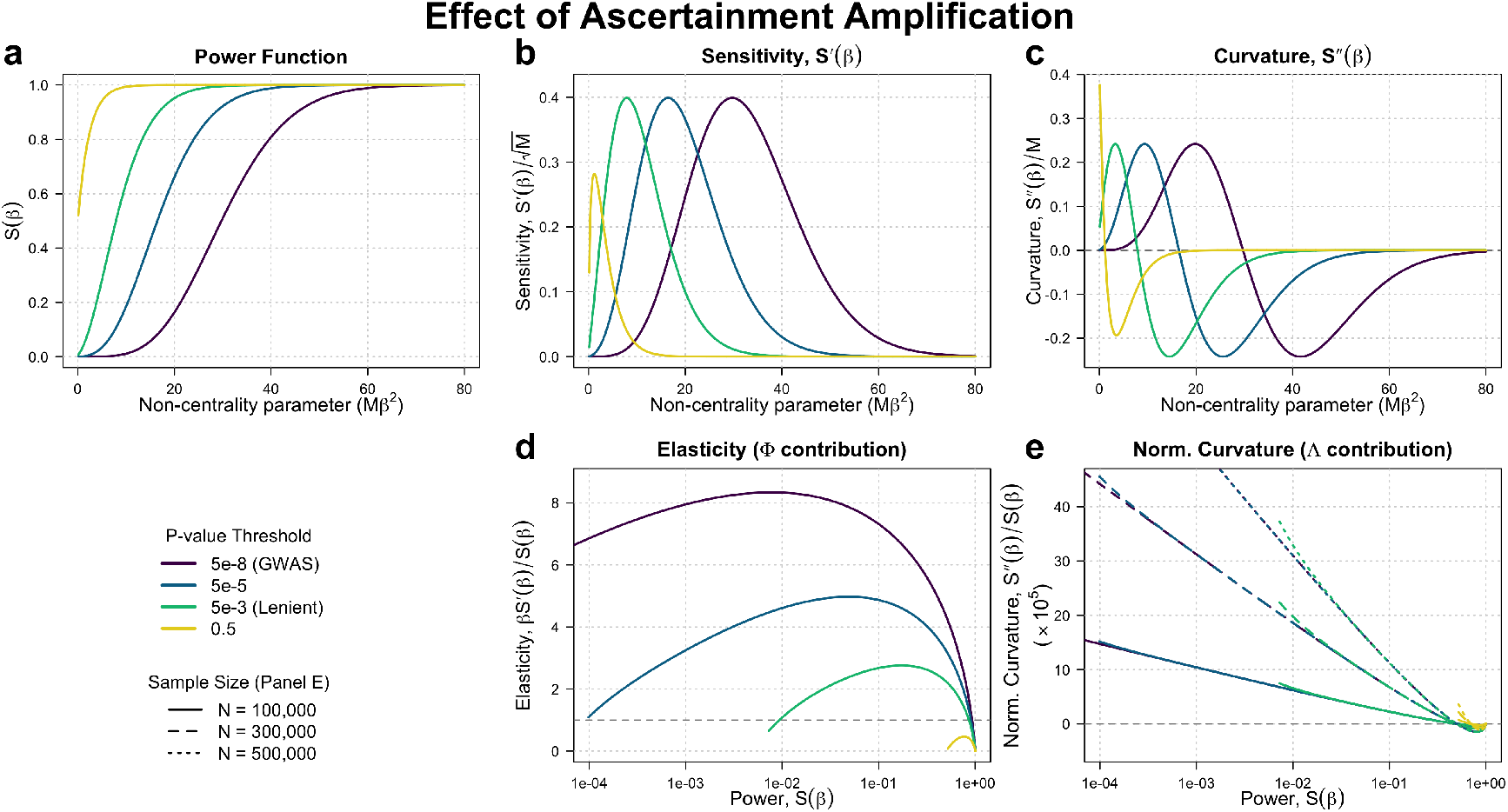
Effect of ascertainment amplification. (a-c) The baseline ascertainment probability (power) function, *S*(*β*), and its first and second derivatives, plotted as a function of the *χ*^2^ non-centrality parameter of the GWAS test statistic distribution (*Mβ*^2^) for four different p-value thresholds. The sensitivity *S*^′^(*β*) (b) and curvature *S*^′′^(*β*) (c) are scaled by 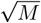 and *M*, respectively, to make the shapes invariant to the GWAS sample size *M* . (d) The elasticity of ascertainment, *βS*^′^(*β*)*/S*(*β*), plotted against statistical power. This quantity (which determines the directional amplification factor Φ) is invariant to sample size but highly sensitive to the significance threshold. (e) The normalized curvature, *S*^′′^(*β*)*/S*(*β*), plotted against power for three different sample sizes (*M* = 100*k*, 300*k*, 500*k*; indicated by line type). Unlike elasticity, normalized curvature is impacted by sample size. This indicates that for a given variant power, larger studies (dotted lines) are significantly more susceptible to the background inflation of SNP counts (Λ) due to a given magnitude of stratification variance.

Here 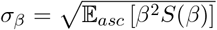 is the standard deviation of effects among ascertained sites. The term 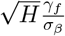 represents the expected bias contribution at a random site included in the PGS, before accounting for ascertainment biases, while 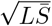 accounts for the accumulation of this signal across the expected number of ascertained variants given the study parameters. Together, these terms quantify the naive bias prior to accounting for ascertainment effects [3].

This naive bias is magnified by the directional ascertainment amplification factor,

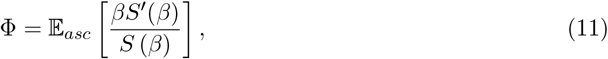

where ^*βS*′ (*β*)^*/S*(*β*) captures the elasticity of ascertainment (i.e. the percentage increase in ascertainment probability resulting from a 1% increase in effect size; see Supplementary Text S1.2.4). This term captures the asymmetry of ascertainment: sites where the causal effect and the bias align (*β*_*ℓ*_*δ*_*f,ℓ*_ *>* 0) are preferentially ascertained, creating a spurious correlation between causal effects and genotype variation along the target axis in the set of ascertained loci. The magnitude of this amplification is determined by the statistical power of the study. Because the elasticity of ascertainment, ^*βS′* (*β*)^*/S*(*β*) is generally greatest for the variants with low or intermediate power (Figure 1d), PGS that include many variants in this regime are susceptible to significant ascertainment driven amplification effects (Φ ≫ 0). Conversely, we would expect PGS dominated by well-powered variants to have have negligible directional amplification effects (Φ → 0).

The second ascertainment factor, Λ, accounts for the global inflation of the SNP count driven by the total stratification load. We express this factor as an expectation over the distribution of ascertained variants,

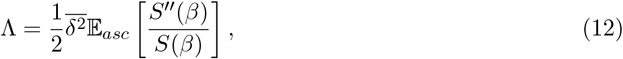

where ^*S*′′ (*β*)^*/S*(*β*) is the normalized curvature of the ascertainment function. Notably, the normalized curvature is large and positive for variants with low statistical power (Figure 1e). Because the power curve is convex in this regime, the increased variance in effect size estimates caused by stratification pushes more variants across the inclusion threshold than it removes. The magnitude of this inflation is scaled by 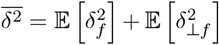, the average total squared confounding bias. Assuming the focal axis represents only a small fraction of total population structure, we expect 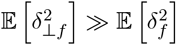, so that this effect is driven primarily by background stratification orthogonal to the target axis. This general mechanism is well-known in statistical genetics as the source of false positive associations [34, 35] and the inflation of heritability estimates [36, 37]. In the context of the PGS test, this effect inflates both the cumulative directional bias (by a factor of 1 + Λ) and the null standard deviation (by a factor of 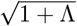) simply by increasing the number of SNPs included, resulting in a net amplification of the naive estimation bias.

### 2.5 Statistical Control via PCA

Standard GWAS methods aim to control for confounding by including covariates, such as the top principal components (**U**). Including these covariates modifies the estimation of effect sizes; geometrically, this is equivalent to projecting the phenotype vector onto the subspace orthogonal to the PCs. We model the corrected effect size estimate as,

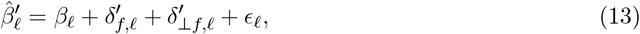

where 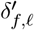 and 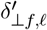 represent the residual bias along the focal and background axes, respectively, after projection onto the orthogonal complement of **U**.

We define the residual target axis *f*^′^ as the projection of *f* onto this subspace,

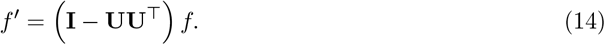

Parallel to the definition of *H*, the residual susceptibility *H*^′^ is the total variance in the GWAS panel genotypes explained by this residual axis,

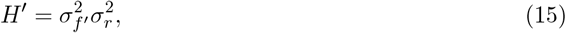

where 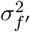 is the variance in the residual target axis *f*^′^, again defined relative to the variance in the prediction panel. Applying the same logic as above to the corrected estimator (see Supplementary Information S1.3), the expected standardized bias becomes under the null,

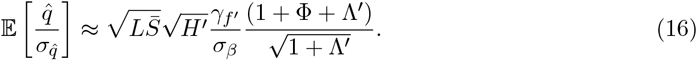

Here, *γ*_*f*_′ represents the magnitude of environmental confounding aligned with the standardized residual target axis 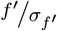 (i.e., the strength of confounding associated with one standard deviation of residual ancestry variation; see Supplementary Text S1.3 and Discussion). The term 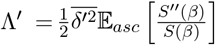 represents the residual inflation of the SNP count attributable to structure not captured by the PCs, where 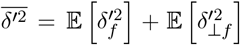 is the residual total confounding. In turn, the directional amplification factor, Φ, is a function only of the causal effect distribution and the power of the study, and so is approximately unaffected by PC correction. Thus, even if total residual confounding is well-controlled (i.e. 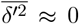), the residual standardized bias will still be large if 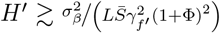. This formalizes the intuition that subtle, axis-specific biases could accumulate to confound polygenic scores even if the total confounding is minimal.

To quantify the effectiveness of PCA we define the statistic *V*_*K*_ as the proportion of the variance in the target axis *f* that is captured by the principal components,

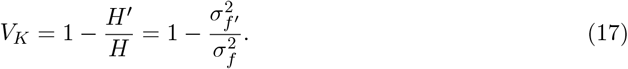

In theory, if enough PCs are included, we would hope that residual susceptibility *H*^′^ = 0, and *V*_*K*_ = 1. In practice, the sample PCs, **Û**, which are actually used to correct for structure are statistical estimates [11**?** ], and so to the extent that they are estimated with noise, we would expect their efficacy to be reduced (i.e. *V* < 1). In the Supporting Information, we provide a generative model linking these factors and a method for estimating *V*. By empirically estimating *V* across a range of scenarios, we can determine whether standard PCA correction is sufficient to mitigate the bias derived above.

## 3 Data

### 3.1 Prediction panel genotype contrasts

Polygenic scores are used in a broad range of applications. Notably, the prediction samples in which they are deployed vary widely in the scale of their genetic structure, and in the extent to which this structure is related to that of the GWAS panel. Our aim was therefore to generate genotype contrasts that span this variation, allowing us to assay the effectiveness of PCA across the range of scenarios in which it is relied upon to prevent confounding. To ensure our framework is robust to the various ways population structure is operationalized, we constructed sets of contrasts using three different rationales, ranging from broad continental groupings to fine-scale geopolitical definitions.

#### Human Genome Diversity Project and Thousand Genomes Project (HGDP1kG)

First, we constructed ten sets of genotype contrasts representing axes of relatively strong human genetic differentiation. We used high-coverage whole genome sequencing data [27] from the 4,094 individuals of the harmonized Human Genome Diversity Project (HGDP) and 1000 Genome Project (1kGP) (hereafter: HGDP1kGP”). We grouped individuals according to the provided `project meta project pop` codes. While these broad labels are imperfect proxies for human genetic variation [38], we chose them as an accessible way to generate axes with varying degrees of differentiation relative to our GWAS panels, thereby testing the limits of stratification control. We selected groups with at least 200 samples, resulting in five groups identified by the consortium descriptors: “East Asia” (eas), “non-Finnish Europe” (nfe), “Africa” (afr), “South Asia” (sas), and “America” (amr). For each of the 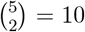 pairwise combinations, we defined a test vector using group membership indicators. These vectors were centered and standardized to unit variance to estimate the genotype contrasts 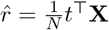 (Figure S2).

Second, we constructed five sets of genotype contrasts along axes of geographic variation for which residual stratification has been a concern in previous studies. These include latitude and longitude within Eurasia, latitude and longitude within non-Finnish Europe, and Sardinia vs. mainland Europe. To construct the Eurasian contrasts, we first constructed a “Eurasian” panel by selecting individuals with “fin”, “nfe”, “eas”, or “sas” `project meta project pop` codes (see Figure S2B and Table S1). We centered and standardized the provided vectors of latitudes and longitudes to have variance one within this panel, and used these standardized test vectors to compute the genotype contrasts. For the non-Finnish European contrasts, we first constructed a non-Finnish European panel by selecting only individuals with “nfe” as their `project meta project pop` code (excluding northern Utah residents with the `project meta project subpop` code “ceu”), and similarly used the provided latitude and longitude as test vectors to generate another pair of genotype contrasts (see Figure S2C and Table S1). To compute the Sardinia vs. Mainland Europe contrast, we constructed a test vector with indicator variables corresponding to individuals from Sardinia (`project meta project subpop` code “sdi”) and all other individuals with the code “nfe” (see Figure S2D and Table S1).

#### British Isles Country of Birth (CoB)

Third, we wanted to investigate axes of relatively subtle variation that were nonetheless well-represented within our GWAS panels. To this end, we selected all individuals from the UKB not belonging to any of our four 100K GWAS panels (see below) whose Country of Birth was listed as Scotland (SCT), England (ENG), Northern Ireland (NI), the Republic of Ireland (RoI), or Wales (WAL), and whose self-identified ethnic background was recorded by UKB code 21000 as `White` (see Table S2 for sample sizes). We then constructed genotype contrasts for all 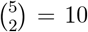 pairs of birth countries in the same manner as above for the HGDP1kGP.

### 3.2 GWAS Panels

We used the UK Biobank [28] to construct six GWAS panels featuring varying levels of genetic diversity and structural alignment with the contrasts described above. First, we used the standard “White British ancestry subset” (WBS) as defined by Bycroft et al. [28], consisting of individuals who self-identified as `White British` and cluster near the centroid in genetic PCA space. This cohort was selected to minimize the amount of ancestral diversity, and thereby minimize the potential for stratification bias. For the HGDP1KG contrasts, we used all genotyped WBS samples (*M* = 408, 625), while for the CoB contrasts we excluded individuals used to compute the test vectors (*M* = 256, 581). Second, we used the entire biobank as a GWAS panel (*M* = 486, 612 for the HGDP1KG contrasts; *M* = 318, 559 for the CoB, with overlap removed).

Third, to investigate scenarios with intermediate levels of diversity, we generated four additional panels (*M* = 100, 000) using a continuous sampling scheme [39]. First, we calculated the median PC 1 and PC 2 coordinates of the full biobank and determined the Euclidean distance of every individual from this centroid. We then randomly sampled 100,000 individuals falling within each of four sampling distances measured in normalized PC units: 5*ϵ*, 10*ϵ*, 50*ϵ*, and 200*ϵ* (where *ϵ* = 10^−4^). Figure 2a visualizes the genetic distribution of these panels alongside the WBS and the full biobank.

**Figure 2:**
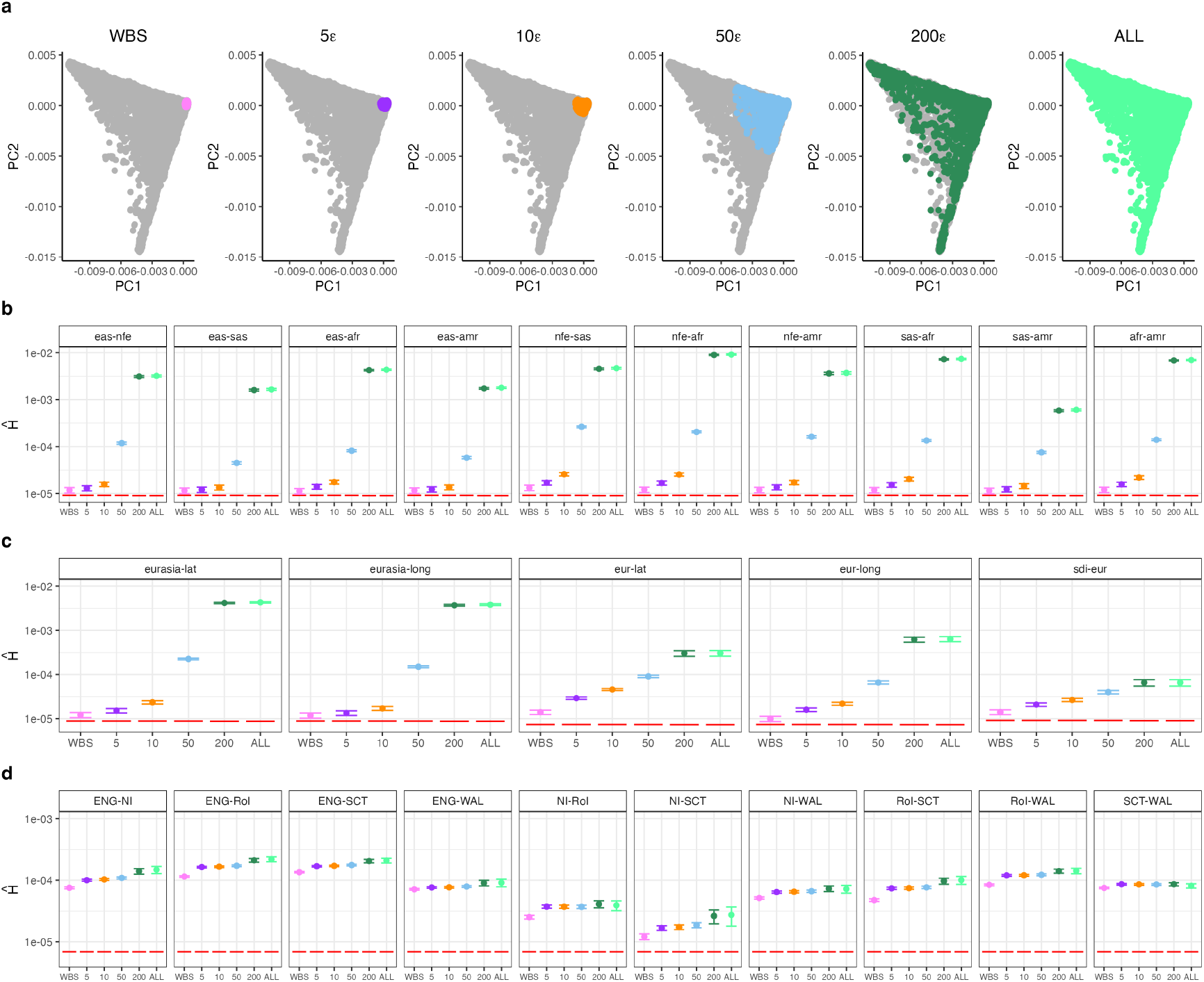
Genetic diversity and susceptibility to stratification across six UK Biobank GWAS panels. (a) Principal component (PC) biplots visualizing the genetic diversity of six distinct GWAS panels constructed from the UK Biobank. The intermediate panels (5*ϵ*, 10*ϵ*, 50*ϵ*, 200*ϵ*) each contain 100,000 individuals randomly sampled from within a specified Euclidean distance from the PC centroid of the full dataset (where *ϵ* = 10^−4^). (b-d) Estimates of the uncorrected susceptibility *Ĥ* (proportion of genetic variance explained by the target axis) for each panel across three sets of contrasts: (b) broad ancestry gradients constructed from geographic labels in HGDP1kGP, (c) continuous geographic gradients within Eurasia and the Sardinia vs. mainland Europe contrast, and (d) country of birth contrasts within the British Isles. Dashed red line indicates the theoretical detection limit (≈ 1*/L*).

## 4 Results

### 4.1 Uncorrected susceptibility to stratification (*H*)

To assess the baseline susceptibility of different study designs to stratification bias, we first estimated *H*—the proportion of genetic variance in the GWAS panel associated with each target ancestry gradient—prior to any statistical correction. We tested the null hypothesis that *H* ≈ 0 (specifically, that the target axis explains no more variance than expected by chance, i.e., *Ĥ >* ^1^/*L*; see Methods section 6.2). After Bonferroni correction, we rejected the null for all 25 contrasts across all six GWAS panels (*p* < ^0.05^/(25×6) = 3.33 × 10^−4^), confirming that some degree of stratification risk exists in every scenario that we tested.

However, the magnitude of this risk varies dramatically depending on the choice of GWAS panel. Unsurprisingly, the largest values of *Ĥ* occur in the widely sampled ALL panel (Figure 2b,c), where contrasts between deeply diverged HGDP1KG samples are associated with as much as 1% of the total genetic variance (*Ĥ* ≈ 0.01). The smallest values, in turn, arise from the more homogeneous WBS panel, where these same contrasts explain orders of magnitude less variance (*Ĥ* ≈ 10^−5^), only slightly above the null expectation.

For the finer-scale British Isles (CoB) contrasts, the pattern is different. The proportion of variance explained is generally intermediate between the extremes seen with the HGDP1KG contrasts, and much more stable across all panels (Figure 2d). However, it is nonetheless the case that *Ĥ* increases with increased sampling breadth in almost all cases, presumably because most individuals in the UKBB who are not included in the WBS have ancestries very closely related to those that are. The fact that *Ĥ* more or less uniformly increases with sampling breadth is thus a particular feature of the UK Biobank and the sampling schemes that we have used, and not a necessary consequence of increasing breadth in general.

### 4.2 Efficacy of Principal Components in capturing target axes

Given that *H >* 0 in all tested cases, we next evaluated whether standard controls for population structure—specifically Principal Component Analysis (PCA)—successfully capture these target axes. We estimated 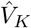, the fraction of the variance along the theoretical target axis *f* that is captured by the top *K* principal components (see theory section 2.5 and methods section 6.4 for details). We evaluated three PC configurations to assess the relative utility of common versus rare variation: 1) 40 common variant PCs, 2) 40 rare variant PCs (*MAF* < 0.01), and 3) the combination of both.

We found a strong relationship between the estimated strength of the stratification signal, *Ĥ*, and the efficacy of common variant PCA (Figure 3a). For target axes with *Ĥ >* 10^−3^ (i.e., HGDP1KG contrasts in diverse panels), 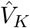 was uniformly very close to 1, while for those with *Ĥ >* 10^−4^, 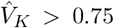 in most cases. Interestingly, the one exception to this pattern was for rare variant PCs used in isolation, which had poor performance specifically for signals of intermediate strength (*Ĥ* ≈ 10^−4^) in most diverse panels (Figure 3a, middle row).

**Figure 3:**
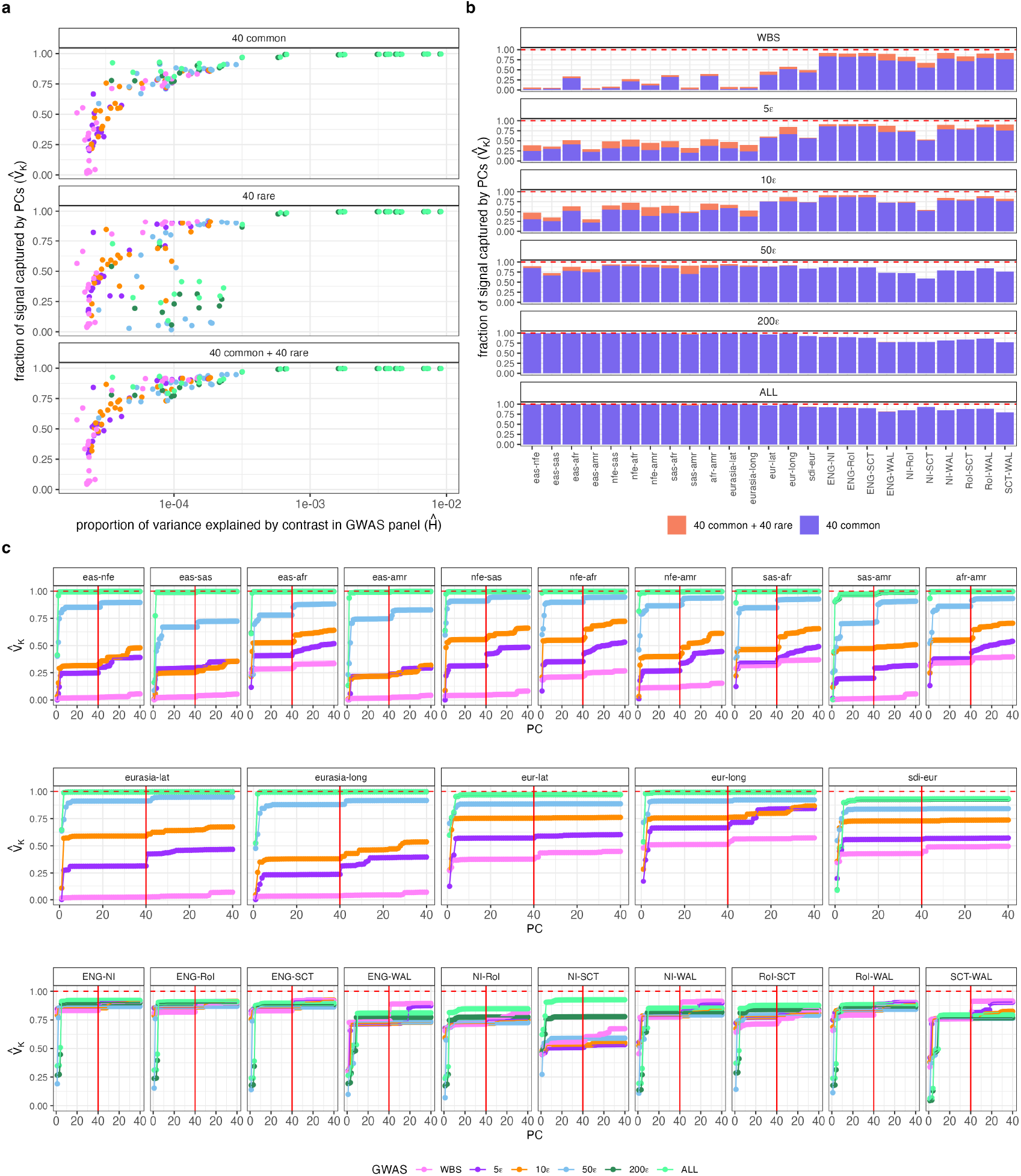
Proportion of variance in target axes captured by common and rare variant PCs. (a) We estimated 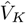 for every GWAS panel/contrasts pair using the top 40 common variant PCs (top row), the top 40 rare variant PCs (middle row) and both types of PCs together (bottom row). In general, the larger the proportion of variance explained by target axis, the larger the fraction of signal captured by PCs. (b) Estimates of 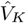 for all GWAS panel/contrast pairs. Common variant PCs explain most of the detectable variance in the target axes with rare variant PCs explaining some amount of additional variance in narrowly sampled panels. (c,d,e) 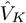 for each common variant PC *K* = 1 to *K* = 40 (left of the red line) and all 40 common variant PCs plus *K* = 1 to *K* = 40 rare variant PCs (right of the red line).

For target axes that explain less variance (i.e. *Ĥ* < 10^−4^), common variant PCs capture a much smaller fraction of the signal. Most of the contrasts in this category are those from the HGDP1KG when analyzed in the narrowly sampled panels (i.e., WBS, 5*ϵ*, 10*ϵ*; see Figure 3b). In these contexts, the inclusion of 40 rare variant PCs alongside common PCs resulted in a measurable increase in 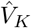. The same was true for some of the CoB contrasts in those same narrowly sampled panels. While this suggests that rare variants capture additional structure missed by common variants in homogeneous samples, the improvement is generally incremental. In no case does the addition of rare PCs transform a contrast for which *V*_*K*_ < 1 into one with *V*_*K*_ ≈ 1.

Finally, in Figure 3c we plot 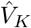 cumulatively, first for the top *K* = 1 − 40 common variant PCs, and then for the top *K* = 1 − 40 rare variant PCs conditional on the inclusion of the common set. Notably, for the GWAS datasets and contrasts that we test, any target axis variation captured by common variants is typically explained by the top ∼ 10 PCs, with components beyond that point adding little. This suggests that common variant PCs beyond the 10^*th*^ either explain so little genetic variation that they are poorly estimated, or that the specific axes they capture are not well-represented within our contrasts. Alternatively, where rare variant PCs do capture significant additional variation, the contributions are often distributed across higher-numbered components or throughout the full set of 40. This lack of a plateau suggests that for at least some of our contrasts, rare variant PCs beyond the 40^*th*^ would likely capture additional variance.

### 4.3 Susceptibility to confounding after population structure correction

While our exploration of the 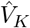 statistics provides insight into the relative utility of different PC sets, the ultimate metric of stratification risk is *H*^′^, the absolute residual genetic variance along the target axis. To assess the limits of standard stratification control, we computed *Ĥ* ^′^ for the most comprehensive correction model (40 common and 40 rare variant PCs). We then conducted the same one-tailed hypothesis test for *Ĥ* ^′^ *>* 1*/L* as described in Section 4.1. After Bonferroni correction, we failed to reject the null hypothesis of zero residual structure for 47 out of 150 GWAS panel-genotype contrast pairs (*p* < 3.33 × 10^−4^), indicating that for these pairings, the residual genetic variation along the target axis is not statistically distinguishable from sampling noise (Figure 4a). While we formally reject the null for the remaining 103 pairs, the estimated values of *Ĥ* ^′^ occupy a narrow range (5.25 × 10^−6^ < *Ĥ* ^′^ < 1.55 × 10^−5^), close to the theoretical limit of detection imposed by sampling noise (1*/L*).

**Figure 4:**
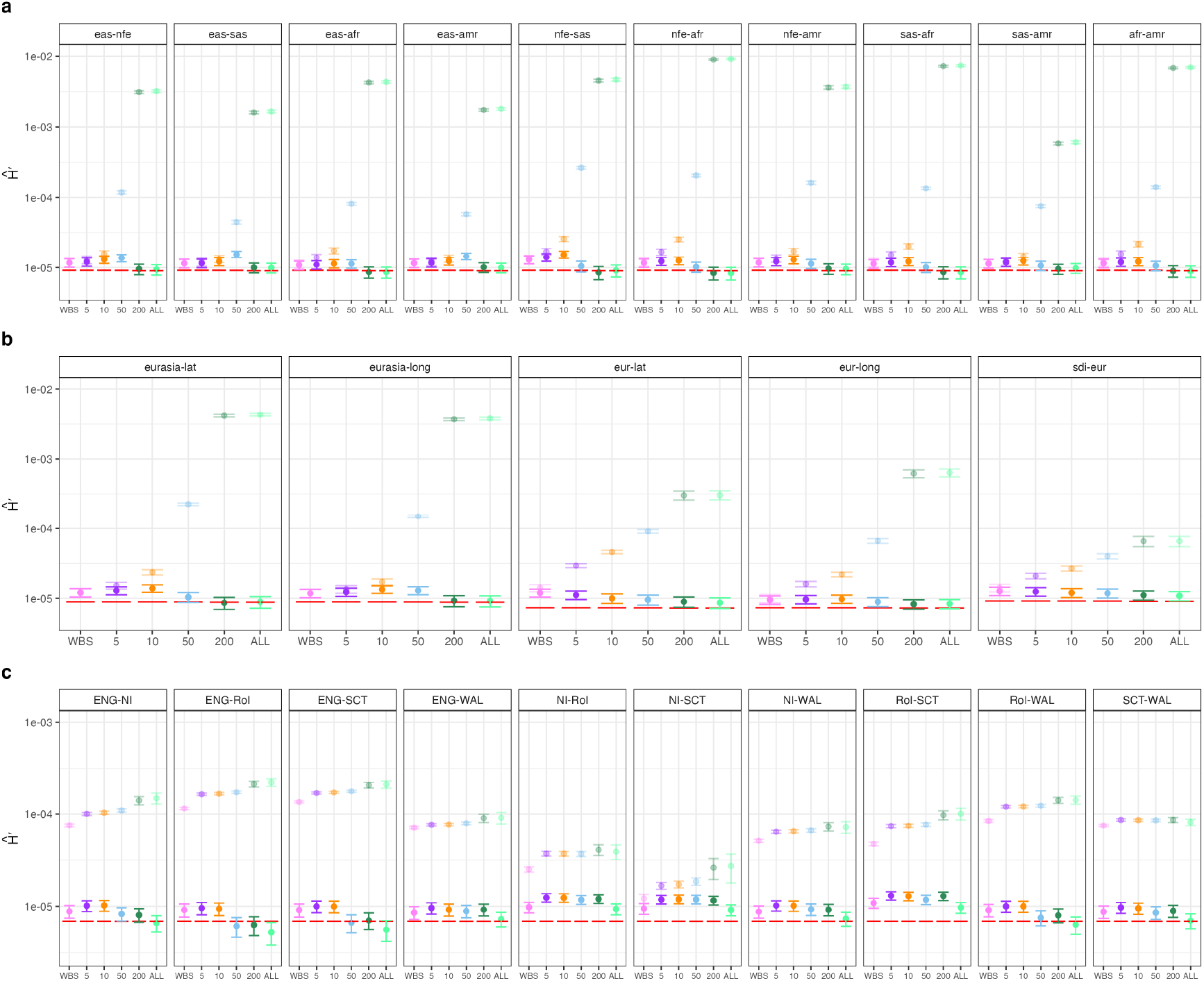
Population structure correction flattens the disparity in stratification risk between diverse and homogeneous panels. We estimated the residual susceptibility *Ĥ* ^′^ after correcting for 40 common variant and 40 rare variant principal components (Methods section 6.2). Estimates are shown for each of the six GWAS panels (x-axis) across three sets of ancestry contrasts: (a) broad pairwise HGDP1kGP gradients, (b) continuous Eurasian geographic gradients (plus Sardinia vs. the mainland), and (c) British Isles country of birth contrasts. For comparison, the uncorrected *Ĥ* values are shown as semi-transparent points. The horizontal red dashed line indicates the theoretical limit of detection ( ≈ 1*/L*). Notably, for many contrasts in the diverse ALL panel, robust correction reduces *Ĥ* ^′^ to levels indistinguishable from the noise floor, comparable to or lower than the residual risk in the restricted WBS.

Nevertheless, examining the correlates of this residual variation reveals interesting trade-offs in study design. We find that PCA is generally more effective at protecting a given axis in a diverse panel than in a homogeneous one. For example, we fail to reject the null for 21 out of 25 contrasts in the diverse ALL panel, compared to 17 in the 200*ϵ* panel, 9 in the 50*ϵ* panel, and 0 in the narrower 10*ϵ*, 5*ϵ* and WBS panels (Figure S9). Thus, while restricting analysis to the WBS reduces the initial uncorrected risk (*H*), using the full diversity of the Biobank allows for more accurate PC estimation, which ultimately results in lower residual susceptibility (*H*^′^).

Finally, we note the independent impact of sample size (*M*). Theory predicts that increasing sample size improves the accuracy of PC estimation, thereby reducing *H*^′^ [11**?** ]. This effect is evident when comparing the 200*ϵ* panel (*M* = 100, 000) to the ALL panel (*M* ≈ 318*k*, or 486*k* depending on the contrast). Despite having nearly identical levels of uncorrected susceptibility (*H*), PCA often performs better in the larger ALL panel, particularly for contrasts where the initial signal is weak. Across all pairs, the product *M Ĥ* (a rough measure of the amount of information available to estimate the structure [40, 41**?** ]) explains a small but statistically significant portion of the variation in the residual variance *H*^′^ (*R*^2^ = 0.056; *p* = 0.0035; Figure S10).

### 4.4 Polygenic score association test results from 17 phenotypes

#### 4.4.1 Global patterns of bias and ascertainment effects

Theoretically, we expect the bias in uncorrected polygenic score association tests to increase with the uncorrected susceptibility *H* (or residual susceptibility *H*^′^), with the slope determined by the number of SNPs included and the magnitude of confounding relative to the causal genetic effect (Equations 10 and 16; Figure 5a). To evaluate this expectation empirically, we conducted polygenic score association tests for 17 UKBB phenotypes using our 25 sets of genotype contrasts and two GWAS panels (WBS and ALL). For each combination, we compared results from GWAS run with and without correction for both common and rare PCs, with and without an LMM included, using both stringent (*p* < 5 × 10^−8^) and lenient (*p* < 5 × 10^−3^) ascertainment thresholds.

**Figure 5:**
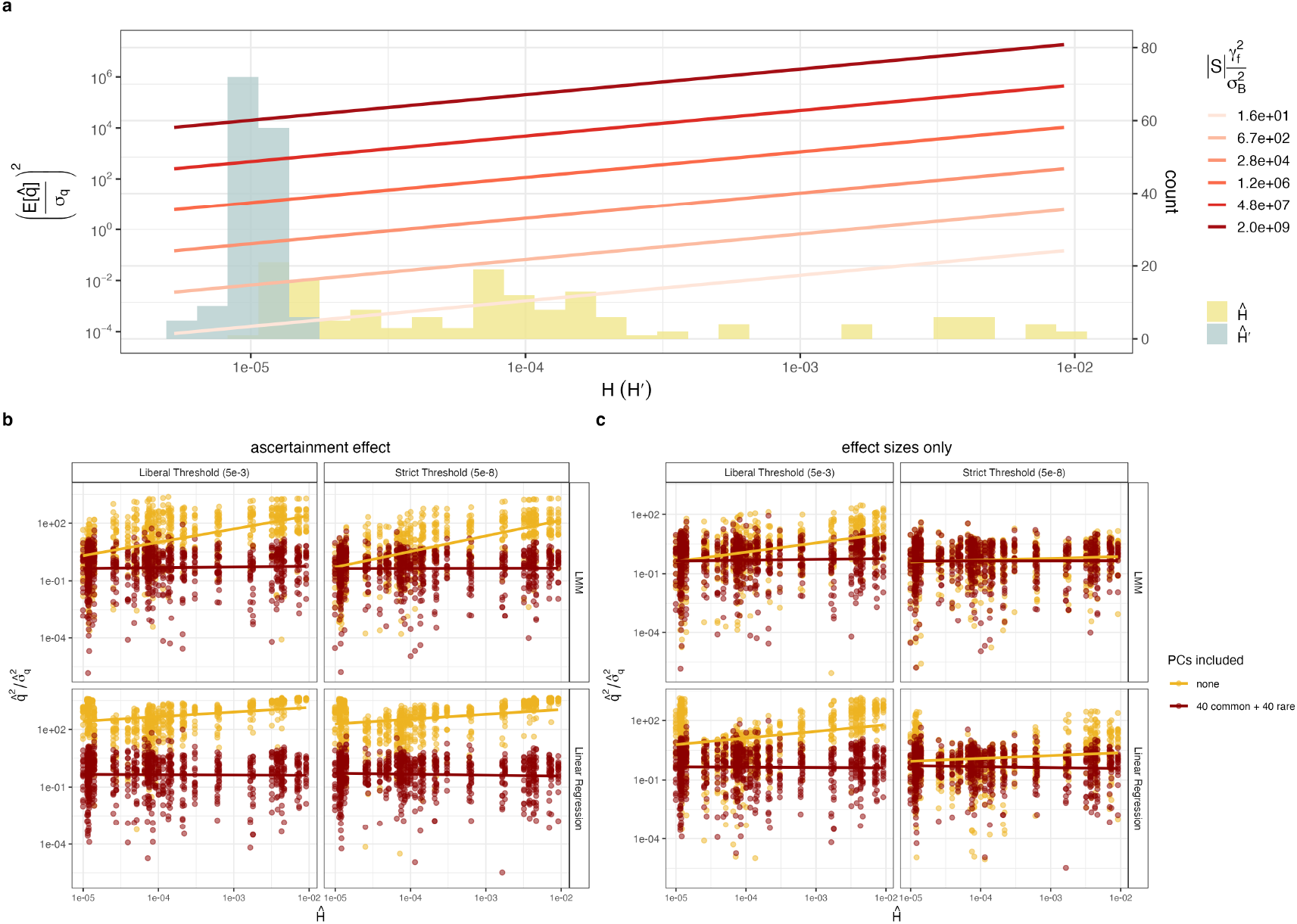
Principal component analysis effectively mitigates stratification bias across diverse study designs. (a) Histograms show the distribution of uncorrected susceptibility *H* (yellow) and residual susceptibility *H*^′^ (blue) across GWAS panel-genotype contrast pairs, while lines show the expected bias as a function of susceptibility. (b, c) Standardized squared test statistics 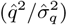 plotted against uncorrected susceptibility (*H*). (b) Standard ascertainment: When SNPs are ascertained using the same model used to estimate effects, uncorrected models (yellow) exhibit significant inflation. This is most severe in simple linear regression (bottom left), driven by the compounding effects of high ascertainment elasticity (Φ) and SNP count inflation (Λ). However, robust correction (red) effectively breaks this relationship, maintaining test statistics near the null expectation. (c) PC-Corrected ascertainment: To isolate estimation bias from ascertainment bias, SNPs were ascertained using PC-corrected statistics. The substantial reduction in inflation for the uncorrected linear regression compared to (b) confirms the potent role of ascertainment in amplifying stratification risk in the absence of correction.

In Figure 5b, we plot the standardized squared test statistics 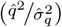 against *H* across all 850 GWAS panel-contrast-phenotype combinations. As expected given our uniformly low estimates for *Ĥ*^′^, including principal components effectively breaks the relationship between stratification susceptibility and bias; for the corrected scores, the test statistics remain uniformly low even for contrasts with high initial *H*.

In the uncorrected cases, however, we observe distinct patterns of inflation that illustrate the theoretical components of ascertainment bias derived in Section S1.2. Under the lenient threshold, the simple linear regression model (Figure 5b, bottom left) exhibits massive inflation even for contrasts with very small values of *H*. This reflects the compounding effects of a large total stratification load (Λ), which dramatically inflates the number of ascertained SNPs (Figure S11), and the high elasticity of the ascertainment function in the low-power regime (Φ), which strongly amplifies the directional bias. In contrast, the uncorrected LMM (Figure 5b, top left) shows significantly less inflation at low levels of *H*. This suggests that the random effect in the LMM successfully dampens the background stratification load, thereby reducing the SNP count (Figure S11) and limiting Λ. However, the LMM fails to completely remove the specific directional confounding aligned with the target axis, which continues to drive bias accumulation as *H* increases. This illustrates how LMMs used in isolation, without the additional flexibility of fixed effect PCs, are insufficient to fully remove residual confounding. These patterns are largely replicated when using the more stringent genome-wide significance threshold. In this regime, the LMM reduces inflation slightly more effectively for low values of *H*, principally by including fewer SNPs (Figure S11), while the behavior of the uncorrected linear regression remains roughly the same.

We confirm these mechanisms in Figure 5c by decoupling effect estimation from ascertainment. Here, we ascertain SNPs using PC-corrected summary statistics (hopefully ensuring that Λ^′^ is small in addition minimizing *H*^′^) but calculate the score using the uncorrected effect size estimates. This procedure dramatically reduces the inflation in the linear model and lowers the slope of the bias in the LMM, confirming that ascertainment bias acts as a potent amplifier of stratification risk in the absence of effective PC correction.

#### 4.4.2 Interpreting significant polygenic score-ancestry associations

Finally, we demonstrate how our framework can help interpret statistically significant associations. Given that we find that the ALL panel has the largest sample size and is protected equally as well or better than all of the other panels, we focus our analysis on this panel. After correcting for multiple testing, we identified seven significant associations between polygenic scores and ancestry gradients in models corrected with an LMM plus 40 common and 40 rare PCs (Figure 6; see Table S4 for all significant results obtained without an LMM and/or in the WBS panel). Notably, the observation that *H*^′^ is statistically indistinguishable from zero does not guarantee that stratification bias is absent; it merely establishes that residual structure is below the limit of detection. Because polygenic scores aggregate effects across many loci, even undetectable levels of residual structure could compound into significant bias if environmental confounding is sufficiently strong.

**Figure 6:**
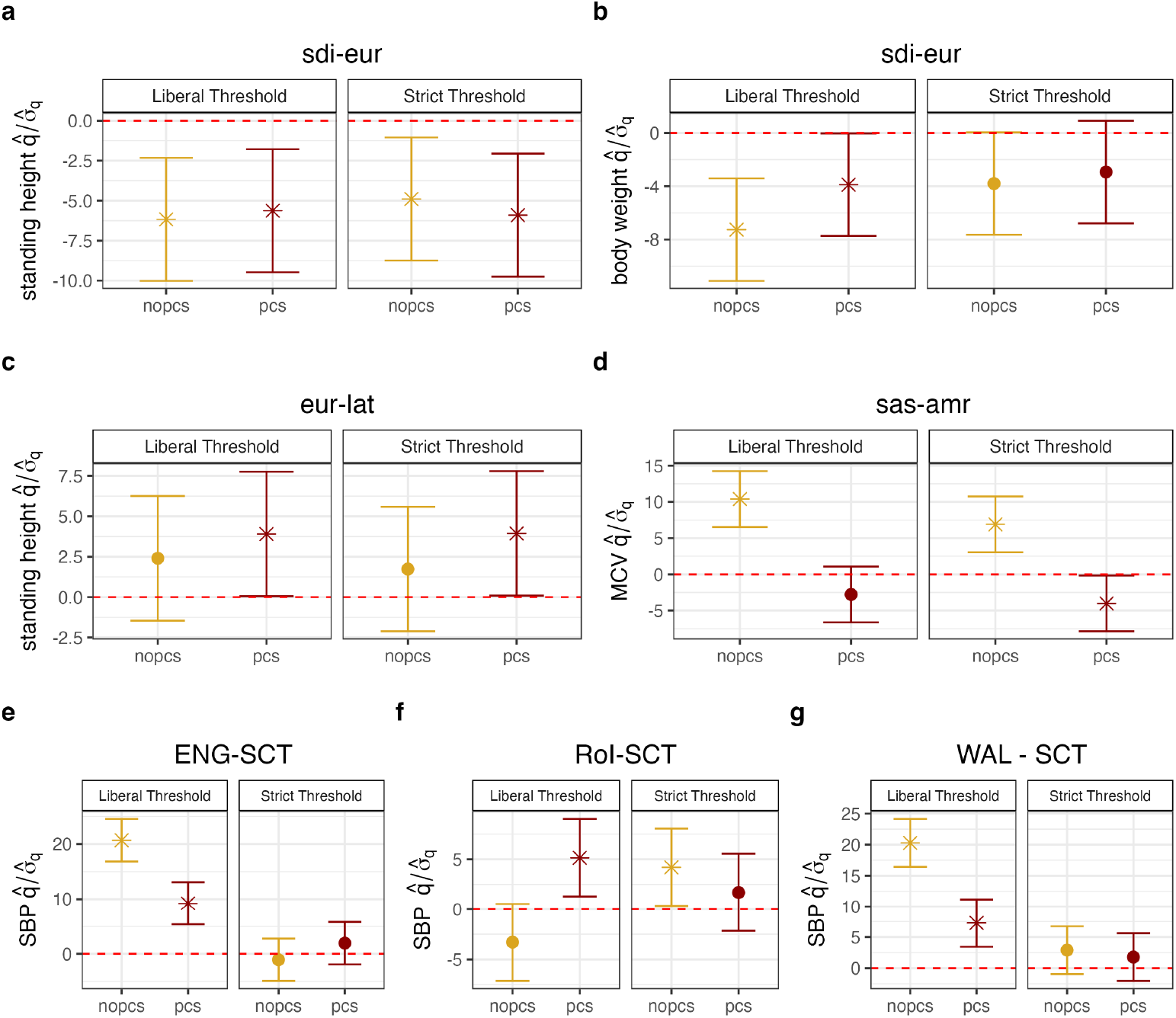
Significant polygenic score-ancestry associations in the ALL panel of the UK Biobank. We conducted polygenic score association tests for 17 phenotypes across 25 ancestry contrasts. (a-g) Standardized test statistics 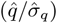 for contrast/phenotype pairs that were significant after Bonferroni correction in at least one PCA-corrected analysis condition. Points represent the test statistic calculated using effect sizes from either uncorrected (yellow) or PC-corrected (red) GWAS models (in all cases including a linear mixed model). Error bars indicate 95% confidence intervals derived from block jackknife standard errors and adjusted for the 17 × 25 = 425 tests performed under each analysis condition. Asterisks (*) denote associations that remain significant after correction for multiple testing (*p* < 0.05/425).

To distinguish potentially robust biological signals from artifacts of residual stratification, we estimated the magnitude of environmental confounding required to generate the observed signal purely as a false positive (Table 2; Methods section 6.6). Because our point estimates of residual susceptibility (*Ĥ* ^′^) are often indistinguishable from zero, we first established a plausible upper bound for the residual stratification risk, 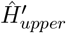, defined as the upper limit of the one-sided 95% confidence interval around a bias-corrected estimate of *Ĥ* ^′^. Using this conservative estimate, we calculated a “naive” critical confounding value 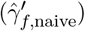 assuming that the bias is driven solely by estimation error (i.e., Φ = 0). To contextualize this magnitude, we also estimated 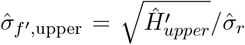, which quantifies the standard deviation of the residual ancestry axis relative to the original population difference defined by the test axis. Together, these metrics imply that to generate the observed signal as a false positive, an environmental variable would need to shift the phenotype by 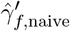 standard deviations per unit of residual ancestry variation along a gradient with a standard deviation that is 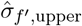 as large as that of the original test axis.

**Table 1:**
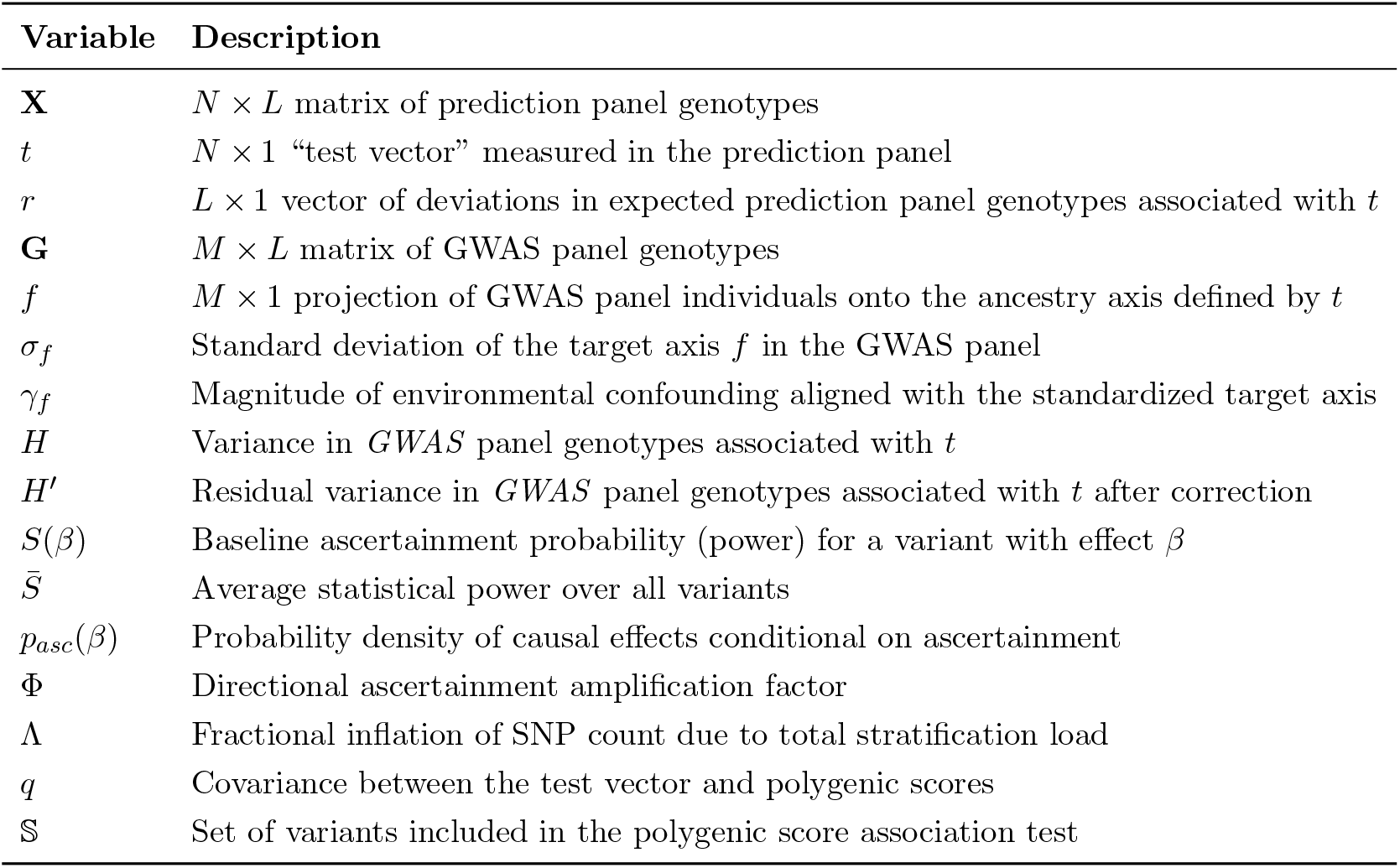
Variable descriptions. Summary of key variables and parameters used in the analysis.

**Table 2:**
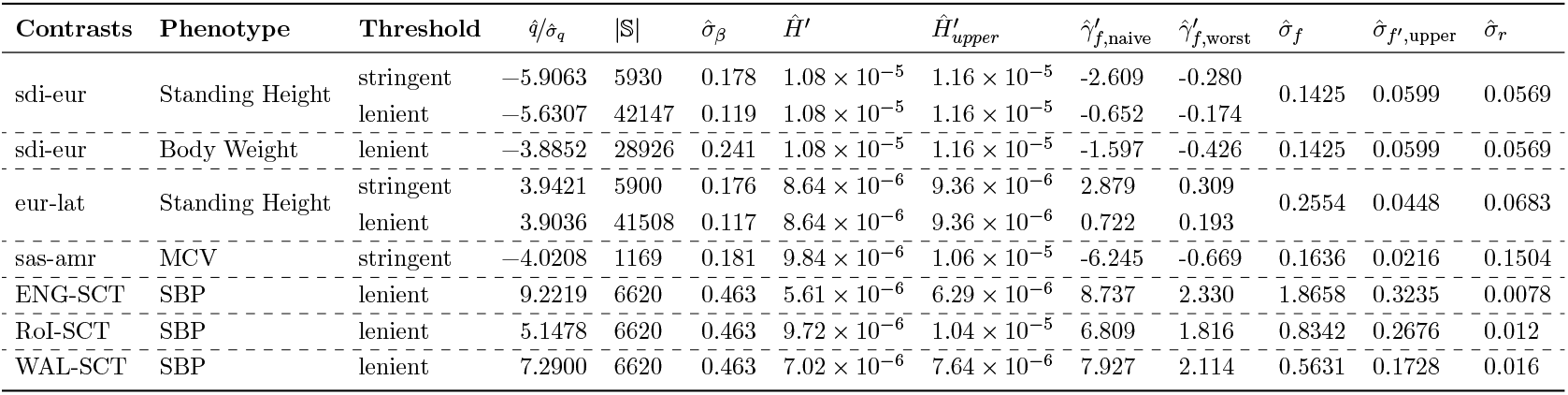
Sensitivity of significant associations to residual confounding. We calculated the critical confounding magnitude 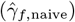 required to explain each significant association in the ALL GWAS panel purely as a stratification artifact. This value represents the required phenotypic shift (in standard deviations) per unit of residual ancestry variation. The naive estimate assumes no ascertainment amplification (Φ = 0), while the worst-case estimate 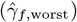 assumes maximal amplification based on theoretical elasticity limits (Φ ≈ 8.33 for stringent, Φ ≈ 2.75 for lenient). To contextualize these magnitudes, we report the estimated standard deviation of the target axis, *f*, before correction 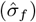 and the upper bound of the standard deviation after correction 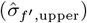, as well as the genotypic standard deviation along the original test axis 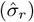. Low absolute values of 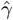 combined with small residual variances 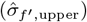 indicate signals that are sensitive to subtle residual confounding.

However, as shown in our theoretical analysis, ascertainment amplifies this bias by a factor of (1 + Φ), meaning the naive critical value is likely an overestimate of robustness. Since the true distribution of effects is unknown, we cannot estimate Φ precisely. To bracket the risk, we calculated a “worst-case” critical value 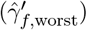 by assuming the score is dominated by variants with maximal ascertainment elasticity (Φ_max_). Based on the theoretical limits derived in Supplementary Section S1.2.4, we scaled the naive estimates by (1 + Φ_max_), using Φ_max_ ≈ 8.33 for the stringent threshold and Φ_max_ ≈ 2.75 for the lenient threshold.

These diagnostics reveal distinct risk profiles among the significant results. For the well-studied divergence in height polygenic scores between Sardinian and mainland European samples [29, 30, 42, 43], the signal appears fragile under a lenient threshold 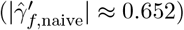. While the stringent threshold raises the naive barrier 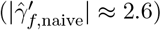 by reducing the number of included variants and increasing their effect sizes, it also increases the amplification factor, dropping the worst-case critical value back to a fragile range 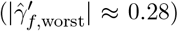. This suggests that an environmental confounder with a modest correlation to the residual ancestry axis could explain the result. However, this interpretation must be calibrated by the scale of the residual variance itself. For the Sardinia contrast, the upper bound of the residual standard deviation is only 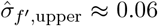. This means that to generate the observed signal as a false positive, the environmental variable would need to shift the phenotype by ≈ 0.3 standard deviations across an ancestry gradient that is only 6% as strong as the original ancestry difference between the Sardinian and mainland European samples. While not impossible, this requirement for non-trivial environmental sensitivity to a very subtle genetic gradient strengthens the case that the signal is not purely an artifact. We observe a similar pattern for the divergence in body weight PGS between the Sardinian and mainland European samples, and for the difference in the MCV PGS between the AMR and SAS samples. We also note that our eur-lat contrast includes the Sardinian sample, so we cannot be sure that the eur-lat latitudinal signals, which also have similar patterns, are completely independent of the Sardinia vs. mainland signal.

In contrast, the novel signals of divergence in Systolic Blood Pressure (SBP) between the SCT sample and the WAL, RoI, and ENG samples appear robust for a slightly different reason. For these associations, the upper bound of the residual ancestry standard deviation is notably larger 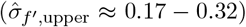. This occurs because these fine-scale contrasts correspond to very subtle test panel ancestry gradients (i.e. they have a small 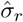). As a result, even though our rough upper bound on the residual genetic variance, 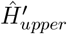, is similar across all contrasts, it represents a larger fraction of the original signal in the test panel. This highlights an important limitation: for subtle ancestry gradients, we can have less confidence about how well PCA has reduced the variance along the target axis relative to that of the externally defined test vector. However, despite this uncertainty, the confounding magnitude required to explain the signal is implausibly large 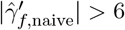 in all cases). Even under the worst-case ascertainment scenario, the confounding required to explain these signals remains high 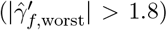, suggesting that these differences are unlikely to be driven by uncorrected population structure confounding as modeled here, offering a high-confidence target for future investigation.

## 5 Discussion

The promise of polygenic scores lies in their ability to predict phenotypes from genetic information alone. However, this promise is compromised when the prediction relies on correlations confounded by population structure. To address this, we developed a framework to quantify the susceptibility of polygenic scores to stratification bias along specific, externally-defined ancestry gradients. By defining the uncorrected susceptibility (*H*), the residual susceptibility after correction (*H*^′^), and the ascertainment amplification factor (Φ), we provide direct metrics for the risk of stratification that move beyond global diagnostics like the LDSC intercept [37].

Applying this framework to the UK Biobank, we found that the magnitude of uncorrected susceptibility varies dramatically with study design. In diverse panels, broad ancestry gradients explain a substantial proportion of genotypic variance (*H* ≈ 0.01), whereas in homogeneous subsets like the WBS, this variance drops by orders of magnitude. However, our results challenge the assumption that restricting analysis to homogeneous groups is the only, or even the best, strategy for safety [28, 44]. We found that robust statistical correction (using top PCs) in diverse panels reduces residual susceptibility (*H*^′^) to levels comparable to, or even marginally lower than, those achieved by sample restriction. This suggests that the increased power and precision of PC estimation in diverse cohorts can outweigh the benefits of minimizing initial structure. This finding supports a movement toward analyzing diverse cohorts as single units rather than fragmenting them [45], provided that appropriate correction methods are employed.

Our analysis of the efficacy of PCA (*V*_*K*_) illuminates the mechanism behind this trade-off. Common variant PCs efficiently capture deeper structure, often removing *>* 99% of the signal for broader ancestry gradients. In contrast, more subtle axes of structure are generally not captured as well by common variants. In these cases, we found that adding rare variant PCs alongside common variant PCs provided some additional protection, though the improvement was modest. However, we expect the utility of rare variants would be greater for even subtler axes of structure [46]. Future studies could leverage our framework to systematically compare the efficacy of different population structure estimators (e.g., fine-scale ancestry components [47] or IBD-based methods [45]) in controlling for confounding across different axes of interest.

However, our theoretical analysis relies on several simplifications, such as the convention of standardizing genotypes to unit variance. This implicitly assumes that all variants share the same relationship to population history, whereas in reality, lower-frequency loci reflect more recent demographic events than high-frequency loci [46]. In principle, parameters like *H* and the target axis *f* could be redefined to account for this frequency-dependent structure, which would be particularly valuable for analyzing stratification at very fine geographic scales. Importantly, a key principle of our framework is that the variants used to define the target axis *f* should match the set of variants eligible for the polygenic score. Thus, while we used only common variants (MAF *>* 0.01) to estimate our metrics, this is appropriate because this is the set from which we construct our polygenic scores. Applying these methods to rare variants would thus require that the definitions of structural susceptibility be adjusted accordingly.

Notably, we found that for all tested contrasts, the residual susceptibility after correction, *H*^′^, was close to the theoretical limit of detection (i.e., *Ĥ* ^′^ ≈ 1*/L*). While this indicates successful removal of the bulk of the signal, it also highlights a fundamental limitation: we cannot distinguish “zero residual structure” from “residual structure below the limit of detection.” Because the standardized bias scales with the square root of the number of included SNPs 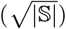 and with the reciprocal of the standard deviation of effects among ascertained sites (1*/σ*_*β*_), even undetectable levels of residual structure can compound into significant bias for polygenic scores that include many sites with very small effect sizes. This implies that “infinitesimal” polygenic scores that aggregate effects across the entire genome are inherently susceptible to very subtle residual stratification biases.

Future work could potentially lower this detection limit or provide assurances beyond it. In this study, we restricted our analyses to a set of high-quality pruned SNPs to minimize the potential for inflation of our *Ĥ* statistic due to linkage disequilibrium (LD). However, future methods could use a larger number of variants (*L*), either by modeling LD to estimate the effective number of independent variants, or by incorporating explicit haplotype-based information to improve resolution [45, 47, 48]. Alternatively, theoretical predictions from Random Matrix Theory regarding the accuracy of PCA [49, 11] could be used to place tighter bounds on residual structure than are currently possible through empirical estimation alone [**?** ].

Regardless of the detection limit, the translation of residual structure into systematic bias is further compounded by the ascertainment process itself. Our theoretical model identifies a directional amplification factor, Φ, which arises because variants aligned with stratification bias are preferentially selected for inclusion in the score. This effect is greatest for variants with low-to-intermediate statistical power. In our empirical analysis, we crudely bracketed this risk by calculating critical values under “naive” (Φ = 0) and “worst-case” (maximal Φ) assumptions. The worst-case value is likely conservative, particularly for the stringent genome-wide significance threshold, where it implicitly assumes that power is on the order of *S* (*β*) ≈ 0.01 for ascertained variants—an implausibly low value for included variants. However, even at substantially higher power, the amplification effect can still be substantial, especially for more stringent thresholds (Figure 1). Thus, while our worst-case bounds establish a lower limit on robustness, the true amplification effect lies somewhere in between. Future studies could refine this bound by using more sophisticated statistical machinery that models both the distribution of causal effects and the power function for ascertained variants, allowing for an explicit estimate of Φ rather than a bracketed range.

To demonstrate the practical utility of this framework, we applied our diagnostic to several significant PGS-ancestry associations. We replicated the well-known divergence in height polygenic scores between Sardinia and mainland Europe [29, 30, 22]. Following the discovery of significant residual confounding in the GWAS meta-analyses that originally supported this result [50, 51], subsequent re-confirmation of the signal has relied on assuming independence between the divergence of mainland Europe vs. Sardinia and the structure found in the evolutionarily diverged Biobank of Japan [43]. While this independence assumption is sensible, we note that we find evidence for very subtle overlaps in population structure even between the WBS and seemingly unrelated contrasts.

To rigorously interpret our analysis of this signal, it is useful to consider a concrete thought experiment regarding the nature of the environmental risk. We might naively suppose that the GWAS panel variation along the target axis *f* arises entirely from the inclusion of individuals sampled in exactly the same way as the Sardinia vs. Mainland test panel, and that the confounding is due solely to a mean environmental difference between these two groups (note that this supposition is clearly at odds with known human demographic history and the sampling of the UK Biobank, but serves as a useful limiting case). Under this assumption, the environmental difference required to explain the signal is scaled by 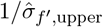. Taking the stringent threshold results as an example, our numbers imply that under naive assumptions about ascertainment, there must be a 2.6/0.06 ≈ 43 standard deviation difference in height between the “Mainland” and “Sardinian” groups in the GWAS panel. Even under worst-case ascertainment assumptions, this number would only fall to 0.28/0.06 ≈ 4.7 standard deviations. Since such massive environmental differences are implausible, we can effectively rule out simple global confounding under the assumptions of this thought experiment.

However, the more realistic risk is that there is some local environmental substructure within the UK Biobank that happens to correlate with the residual axis of confounding left behind by PCA. In this scenario, the risk is determined directly by the residual critical value 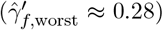. While this value is small enough to be physically possible, it still represents a non-negligible environmental effect that must align perfectly with a residual axis along which individuals vary little in ancestry 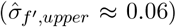 and which explains almost no variance in genotype 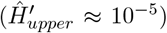. The fact that the signal requires such specific environmental alignment to a subtle genetic feature (together with the independent replication via the Biobank of Japan GWAS) increases our confidence that it is not an artifact. Thus, while the preponderance of the evidence supports the validity of the Sardinia height signal, our analysis demonstrates the difficulty of formally ruling out stratification bias when working with population-based GWAS. For this reason, the use of family-based effect estimates remains the gold standard [52], and the development of methods to integrate family- and population-based effect estimates provides a promising avenue for progress [20].

In comparison, the novel signals of divergence in Systolic Blood Pressure (SBP) between the SCT sample and each of the WAL, RoI, and ENG samples appear more robust. For these associations, the required confounding to explain the signal is implausibly large 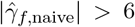 in all cases), suggesting that these differences are unlikely to be artifacts of population structure as modeled here. This result is an interesting target for future study, particularly given historical patterns of cardio metabolic health disparities in Scotland [53].

Finally, we note important limitations of our modeling framework. First, our derivation assumes independent causal sites. In reality, GWAS variants are often tag SNPs in linkage disequilibrium (LD) with true causal variants. However, we expect our core conclusions to hold if the “true causal effect” in our model is replaced by a “tagging effect” (the causal effects of linked variants convolved with their LD to the focal variant). The ascertainment amplification effect (Φ) should still operate on these tagging effects, as tag SNPs aligned with stratification will still be preferentially ascertained. This logic also extends to more sophisticated PGS methods that re-weight variants based on LD [54, 55, 56]. While these methods weight variants differently than the standard thresholding method that we analyze, they ultimately rely on the same summary statistics and are likely subject to similar ascertainment biases. Second, our framework models only stratification driven by drift, gene flow and environmental variation. It does not account for other sources of bias such as participation bias [57], assortative mating [58] or dynastic effects [59, 60].

In summary, our framework provides a path to conducting more rigorous polygenic score analyses by placing quantitative bounds on the plausibility of stratification bias. While no method can guarantee total immunity in population-based GWAS, reporting metrics like *Ĥ* ^′^ and 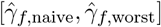 can and provide a transparent accounting of residual stratification risk in PGS-ancestry associations.

## 6 Materials and Methods

### 6.1 Genotype contrasts

#### Human Genome Diversity Project and Thousand Genomes project

We downloaded VCF files from the combined Human Genome Diversity Project and Thousand Genomes Project dataset [27] and converted them to plink2 format [61] after lifting coordinates to the hg19 genome build. As described in Section 3.1, we constructed 15 sets of allele frequency contrasts. For each set, we selected SNPs with greater than 1% MAF in both the test panel and the GWAS panel. We defined the test vector *t* based on population labels or geographic coordinates, mean centered and standardized it such that *Var*(*t*) = 1. We then computed the allele frequency contrast vector 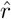 using plink2:

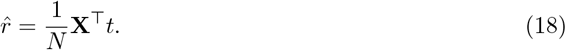

For pairwise contrasts, *t* was coded as {−1, 1} depending on the population label.

#### British Isles country of birth

We selected all individuals from the UKB not belonging to any of our four 100K GWAS panels whose country of birth was listed as Scotland, England, Northern Ireland, the Republic of Ireland, or Wales (see Table S2). We selected SNPs with greater than 1% MAF in both the test panel and the GWAS panel and computed pairwise contrasts as above.

### 6.2 Estimating uncorrected susceptibility (*Ĥ*) and the target axis 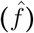

To quantify susceptibility to confounding, we estimate 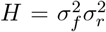. We first estimate 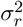 from the prediction panel using the *L* SNPs included in the analysis:

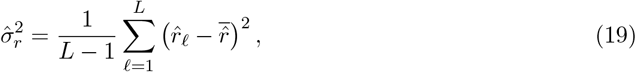

where 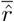 is the mean value. To minimize potential inflation due to linkage disequilibrium (LD), we restricted this estimation to the intersection of SNPs used to compute genetic PCs in the UKB (pruned to pairwise *r*^2^ < 0.1, see [28]) and those for which we had contrast values (*L* = 109, 180 − 145, 247, depending on the contrast).

We used the scoring function `--sscore` in plink2, to estimate the position of each GWAS individual on the target axis:

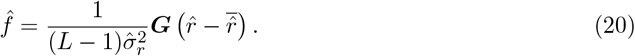

The variance of individual position on this axis is then given by

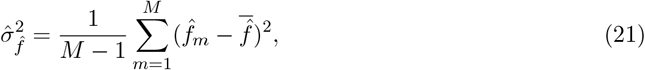

and an estimator of *H* is then:

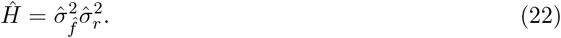

#### Sampling distribution and significance testing

Under the null hypothesis 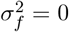, the expectation of *Ĥ* depends on residual linkage. If *ρ* represents the average squared correlation between sites, 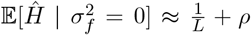. To test significance and estimate standard errors, we used a weighted block jackknife [62] across *B* = 581 approximately independent LD blocks:

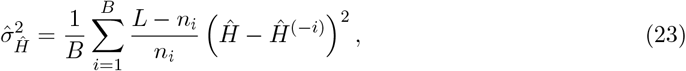

where *n*_*i*_ is the number of SNPs in block *i*, and *Ĥ* ^(−*i*)^ is the estimate omitting block *i*.

In an effort to empirically assess our sensitivity to LD and SNP number, we re-estimated *Ĥ* for the eas-sas contrast, which we might have expected *a priori* to be independent of structure in our narrow GWAS panels, using reduced SNP sets (*L* = 10^3^−10^5^; Figure S3). Decreasing the number of SNPs increases the physical distance between adjacent sites, reducing *ρ* and any potential inflation. However, doing so also increases the sampling noise limit 1*/L*. Thus while reducing *L* eventually eliminates the signal in homogenous panels (WBS, 5*ϵ*, 10*ϵ*), this is precisely the expected pattern for a weak signal. We proceeded by assuming based on our pruning scheme that 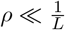.

### 6.3 Population Structure Correction (PCA)

For each GWAS panel, we used the set of 147,604 high-quality, LD-pruned SNPs identified by the UKBB [28] to compute common variant PCs using `--pca approx` in plink2. We extracted the top 40 PCs for use as covariates. For rare variant PCA, we selected variants with *MAF* < 0.01, LD pruned them using `--pca--indep-pairwise` 1000 80 0.1, and subsampled to 1,000,000 variants using `--thin-count` before running PCA and again extracting the top 40 PCs.

### 6.4 Residual susceptibility (*Ĥ* ′) and PCA efficacy 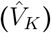

#### Estimating *Ĥ* ^′^

To quantify susceptibility after correction, we regressed the estimated target axis 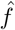 on the top *K* sample PCs (*Û*_*K*_):

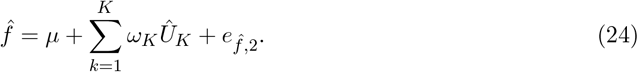

We calculated the fitted residuals, 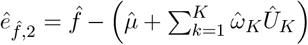, and computed their variance as:

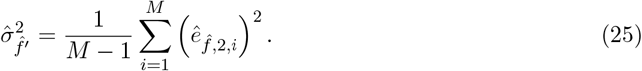

We estimated the residual susceptibility as 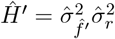, and estimated the sampling variance of *Ĥ* ^′^ using the same block jackknife procedure described above.

#### Estimating 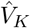 and 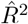

We defined the efficacy of PCA as 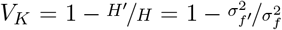. The naive estimator of the target axis variance, 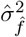, includes estimation noise, which will bias the naive estimator of the ratio statistic 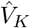. To correct for this, we first estimated the error variance of 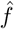 via block bootstrap:

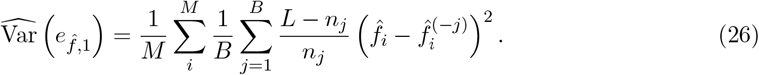

The noise-corrected variance of the target axis is then

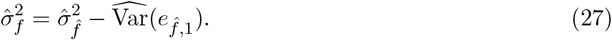

We similarly corrected the residual variance to obtain 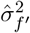 by subtracting the measurement error variance from the variance of the residuals:

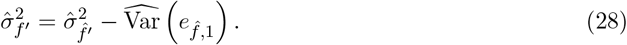

The estimator of *V*_*K*_ is then given by

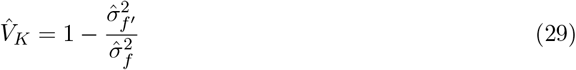

To visualize the relative contributions of signal and error in 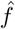, we used these corrected variances to define 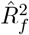, the fraction of variance in 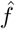 that is due to real underlying variation in ancestry as opposed to estimation noise:

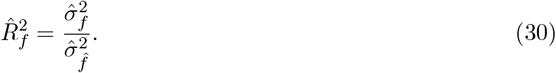

Intuitively, we see that 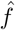 is better estimated (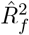 is higher) for pairs with larger values of *Ĥ* than those with smaller values (Figure S4).

Similarly, the fraction of variance in 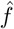 that can be explained by the sample PCs is given by

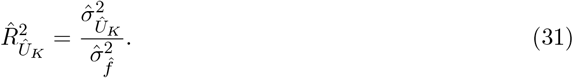

where

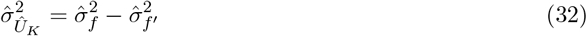

is the variance in *f* captured by the top *K* sample PCs, *Û*_*K*_ .

Notably, 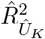 is bounded by 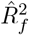 meaning that the PCs cannot explain more variance in 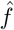 than the fraction due to true variation in ancestry (indeed, the estimator of PC efficacy can be equivalently written as 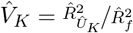. We visualize the relationship between 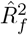 and the cumulative 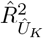 values in Figures S5 and S6.

Finally, to avoid double-dipping—because 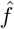 and *Û*_*K*_ are estimated from the same GWAS genotypes—we used a data splitting strategy for 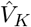 estimation. We computed PCs using SNPs on odd chromosomes and estimated 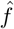 (and its variance components) using SNPs on even chromosomes. We confirmed that 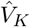 was relatively stable across different chromosome splitting choices as long as both components were estimated from a sufficient number of chromosomes (Figure S8).

### 6.5 GWAS and Polygenic score association tests

We conducted GWAS in the WBS and ALL panels for 17 phenotypes (Table S3) using *regenie* [63]. We ran step 1 of *regenie* we used all directly genotyped variants. In Step 2, we ran two models: 1) a linear mixed model (LMM) using `--pThresh 0.01 --pred --bsize 400`, and 2) a simple linear regression using `--ignore-pred`. Each model was run twice: once with only baseline covariates (age: 21022, sex: 22001, batch: 22000), and once including those covariates plus 40 common and 40 rare variant PCs.

We ascertained SNPs for the PGS using clumping and thresholding in plink2 the clumping parameters `--clump-r2 0.1 --clump-kb 1000` at two significance thresholds (*p* < 5 × 10^−8^ and *p* < 5 × 10^−3^). The raw polygenic score association statistic was computed as 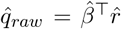. We estimated the standard error, 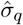, using the block jackknife:

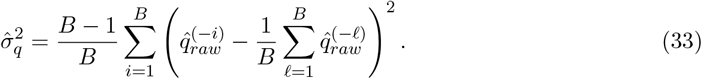

To assess significance, we compared the standardized test statistics 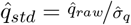 against a Bonferroni-corrected threshold of 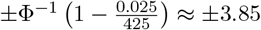, where Φ^−1^ is the inverse of the standard Normal CDF.

### 6.6 Bracketing the critical confounding magnitude

To interpret significant associations, we calculated the magnitude of confounding required to generate the observed signal as a false positive. Because the true bias depends on the unknown ascertainment amplification factor Φ, we bracketed the risk by calculating both a “naive” critical value (assuming Φ = 0) and a “worst-case” critical value (assuming maximal Φ).

First, we computed a bias-corrected point estimate for the residual susceptibility to account for the contribution of sampling noise,

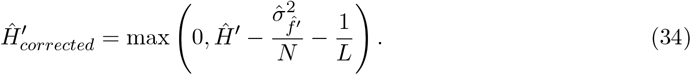

We then determined the upper bound of the one-sided 95% confidence interval,

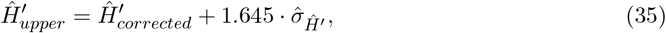

to represent a plausible upper bound on residual amount of genotypic variance associated with the target axis. We then compute a similar plausible upper bound on the residual variance in ancestry along the target axis as

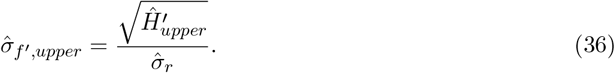

We calculated the naive critical confounding magnitude, 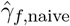, by solving for the confounding strength required to produce a standardized bias equal to our Bonferroni corrected significance threshold (*Z*_*α*_ ≈ 3.851), assuming no ascertainment amplification (Φ = 0):

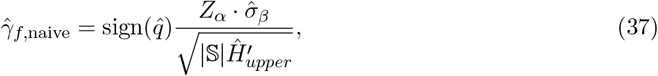

where 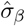 is the standard deviation of effect sizes for the |𝕊| included variants.

Finally, to account for the potential amplification of bias due to ascertainment, we calculated the worst-case critical value,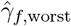. We used the maximum theoretical elasticity values derived in Supplementary Section S1.2.4 (Φ_max_ ≈ 8.33 for the stringent threshold and Φ_max_ ≈ 2.75 for the lenient threshold) to scale the naive estimate:

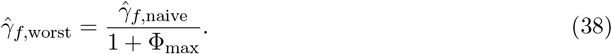

#### Declaration of generative AI and AI-assisted technologies in the writing process

During the preparation of this work the authors used Gemini 2.5 Pro and Gemini 3.0 Pro in order to improve readability and language. After using this tool, the authors reviewed and edited the content as needed and take full responsibility for the content of the publication.

## S1 Supplementary text

### S1.1 The generative model

#### S1.1.1 GWAS panel

To rigorously evaluate the performance of Principal Component Analysis (PCA), we must distinguish between theoretical quantities in the generative model and sample quantities estimated from data. McVean (2009) [64] demonstrated that for any demographic model specifying the expected pairwise coalescent times among individuals, the eigenvectors of the *expected* genetic similarity matrix (∝ 𝔼[**GG**^⊤^]) are functions of those expected times. These eigenvectors define the theoretical PCs, i.e., the axes of variation inherent to the population structure, independent of sampling noise.

Consequently, we can consider a generative specification of the GWAS panel genotype matrix **G** as a realization of these underlying theoretical axes:

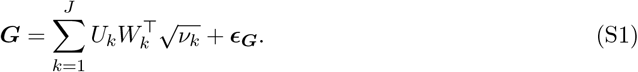

Here, *U*_*k*_ corresponds to the *k*^*th*^ eigenvector of the expected genetic similarity matrix (scaled such that *Var*(*U*_*k*_) = 1), representing the axis of ancestry variation that makes the largest contribution after accounting for the preceding *k* − 1 axes. In turn, *W*_*k*_ is an *L* × 1 vector representing the randomness in allele frequency variation along this ancestry axis (also scaled to *Var*(*W*_*k*_) = 1), 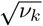 represents the scale of this variation (related to the *k*^*th*^ eigenvalue), and ***ϵ***_***G***_ is the contribution from unstructured random sampling. While we proceed here using principal components, we note that other factor models (such as the structure model, an admixture graph or any other low rank factorization) could fill this role in the generative specification [65].

In the context of this generative model, the target axis *f* and the theoretical PCs *U*_*k*_ can be viewed as alternative rotations of the sample’s shared demographic history; individuals sample their positions on these axes simultaneously via the underlying demographic process. Within this framework, the target axis *f* can be written as a linear combination of the theoretical PCs:

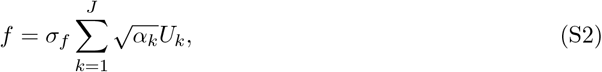

where *α*_*k*_ gives the proportion of variance in *f* that is captured by *U*_*k*_.

Similarly, the vector of allelic deviations *r* lies within the subspace spanned by the variant loadings *W*_*k*_.

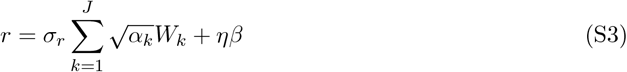

To derive the main text generative model for the GWAS panel, we decompose the theoretical axes *U*_*k*_ by projecting them onto the target axis *f* . We can rewrite Equation S1 as:

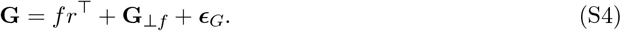

Here, the term *fr*^⊤^ represents the component of genetic variation explicitly structured by the target ancestry gradient. The residual structured variation **G**_⊥*f*_ is composed of the components of the theoretical PCs orthogonal to *f* :

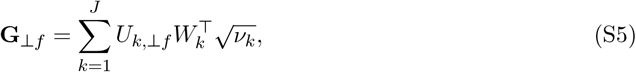

where

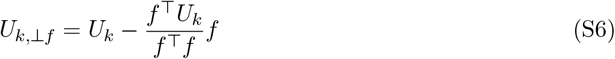

represents the variation in the GWAS panel’s *k* theoretical PC that is independent of the target axis. This matches Equation (3) in the main text.

#### S1.1.2 Prediction panel

We apply a similar generative logic to the prediction panel genotype matrix **X**. Because the prediction panel shares deep demographic history with the GWAS panel, a portion of the variation in prediction panel genotypes is explained by the GWAS panel axes of SNP variation (*W*_*k*_). However, we assume that the prediction panel also contains structured variation that is orthogonal to that of the GWAS panel. We therefore specify **X** as a combination of the GWAS panel theoretical PCs and a set of orthogonal prediction panel-specific axes:

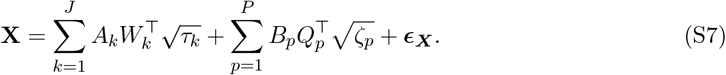

Here, *A*_*k*_ represents the *N* × 1 vector of positions for the prediction panel individuals on the *k*^*th*^ shared theoretical axis (defined by *W*_*k*_). The second summation captures structure specific to the prediction panel: *Q*_*p*_ represents the variant loadings for the *p*^*th*^ orthogonal axis (where *Q*_*p*_ ⊥ *W*_*k*_ for all *k*), and *B*_*p*_ represents the positions of prediction individuals along these axes.

Finally, we consider the generation of the test vector. We assume there exists a latent genetic coordinate *t*^∗^ which is generated by the projection of the prediction individuals onto the shared axes:

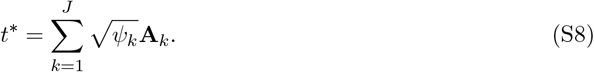

Any structured variation in the test vector that does not project onto the GWAS axes is captured by a second latent variable *t*^†^, defined by the projection onto the prediction panel-specific axes:

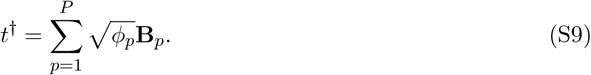

The observed test vector is a realization of these latent states plus measurement error:

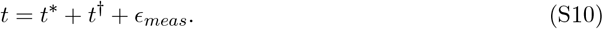

This decomposition allows us to rigorously define the component of the genotype matrix associated with the test vector. We can rewrite Equation S7 as:

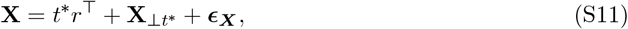

where *r* represents the vector of allelic weights associated with the shared structure *t*^∗^. The residual structured variation **X**_⊥*t*_∗ is composed of the shared variation orthogonal to *t*^∗^ and the variation along the prediction-specific axes:

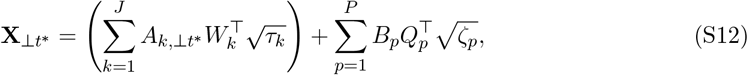

where

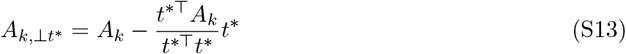

represents the component of the shared axes orthogonal to the test vector. Equation (S11) is identical to equation (2) in the main text.

This specification make the source of the confounding bias clear. While the test vector *t* may be driven by both shared (*ψ*_*k*_) and prediction-specific (*ϕ*_*p*_) structure, only the component associated with the shared axes (*A*_*k*_) projects onto the GWAS panel’s allelic weights (*W*_*k*_). The component associated with the prediction-specific axes (*B*_*p*_) is orthogonal to the GWAS structure (*Q*_*p*_ ⊥ *W*_*k*_) and therefore does not contribute to the allelic weights *r* in the GWAS basis. Thus, the equations for *r* above correctly isolate the component of the test vector that projects onto the shared *GWAS* axes (*W*_*k*_) to generate the confounding signal *f* [3].

### S1.2 Bias in PGS-ancestry associations

The expression for the standardized bias presented in the main text depends on the interaction between stratification and the variant ascertainment process. Here we provide the detailed derivation of this result.

Variants are ascertained for inclusion in a polygenic score based on their estimated effect sizes (e.g., 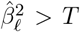, where *T* is a significance threshold). Consequently, the set of included variants, 𝕊, is a random variable that depends on the realized effect estimates. This introduces two distinct sources of ascertainment bias: an inflation in the number of included variants, and a directional bias in their estimated effects. These ascertainment biases in turn amplify the estimation bias. Below, we derive the naive estimation bias, and then show how naive estimation bias is amplified by these ascertainment biases.

#### S1.2.1 Naive estimation bias

We first consider the bias in the absence of any ascertainment effects, assuming the set of included variants 𝕊 is fixed a priori (e.g., all SNPs are included, or SNPs are selected based on an external, unbiased dataset). The estimated PGS covariance is given by

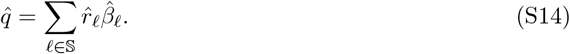

In this scenario, the expected bias depends solely on the systematic error in the effect size estimates 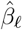. We model the estimated effect as

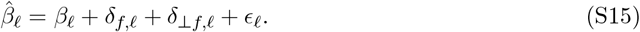

where

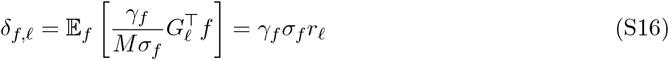

is the bias arising from confounding along the target axis, and

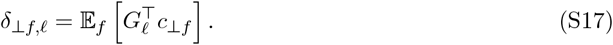

is the bias arising from confounding along orthogonal axes. Because we do not specify an explicit model for *c*_⊥*f*_, we leave the expectation for *δ*_⊥*f,ℓ*_ unevaluated.

The expected bias in the covariance statistic 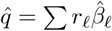 is the sum of the biases across all included loci. Under the null assumption that the causal effects *β*_*ℓ*_ are uncorrelated with the ancestry vector *r*, only the systematic stratification bias *δ*_*f,ℓ*_ contributes

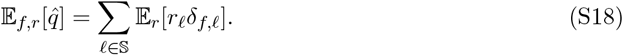

Evaluating the expectation over random variation in *r*, the total bias for a fixed set of size |𝕊| is:

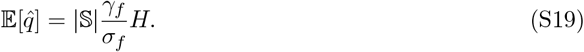

Standardizing by the null standard deviation 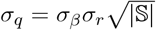, where *σ*_*β*_ is the standard deviation of effect sizes at the sites included in the PGS), we recover the naive estimation bias:

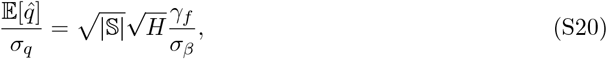

which shows that naively, before accounting for ascertainment effects, we expect the bias to scale with the magnitude of the confounding relative to the effects, the variance explained by the target axis, and the number of SNPs included [3].

#### S1.2.2 Amplification due to ascertainment bias

In practice, the set of included sites 𝕊 is not fixed. To understand the impact of ascertaining variants based on biased estimates, let 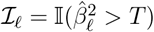 be the indicator variable for ascertainment. The estimated polygenic score covariance with the test vector is explicitly given by the sum over all loci:

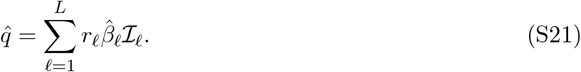

Let *S*(*β*) represent the baseline ascertainment probability, i.e. the statistical power to detect a variant with true effect *β* in the absence of stratification bias (i.e., assuming the estimator follows the distribution 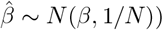.

In the presence of stratification bias *δ*_*f,ℓ*_, the distribution of the estimator shifts to 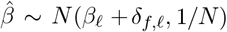. Assuming the magnitude of the bias, *δ*_*f,ℓ*_ is small, We approximate the impact of stratification bias on the ascertainment probability using a first-order Taylor expansion around *δ*_*f,ℓ*_ = 0:

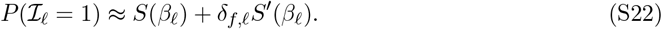

First, we consider the impact on the number of included SNPs. The expected number of SNPs included in the polygenic score is the sum of ascertainment probabilities across all *L* loci. Assuming the bias *δ*_*f,ℓ*_ is independent of the true effects *β*_*ℓ*_ (and thus independent of the sensitivity *S*^′^(*β*_*ℓ*_)), the linear perturbation term sums to zero. Therefore, to a first-order approximation, the expected number of SNPs is simply determined by the average baseline power across the genome, 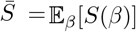:

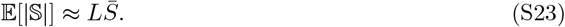

Intuitively, while stratification bias increases the probability of ascertainment at sites where the bias and true effect align (*β*_*ℓ*_*δ*_*f,ℓ*_ *>* 0), it decreases it at sites where they oppose; these effects cancel out in expectation, leaving the total variant count unchanged to first order.

Second, we calculate the directional bias in the covariance estimator 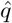. Substituting the expansion into 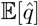, we obtain:

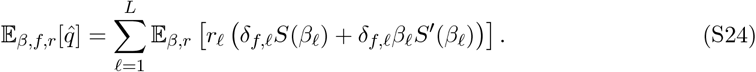

While the opposing effects of stratification cancel out for the total SNP count, they reinforce each other with respect to the directional bias. The sensitivity, *S*^′^(*β*_*ℓ*_), is always positive, so the ascertainment process preferentially selects sites where the causal effect *β*_*ℓ*_ and the bias *δ*_*f,ℓ*_ align (i.e., *β*_*ℓ*_*δ*_*f,ℓ*_ *>* 0), while preferentially excluding sites where they oppose (i.e., *β*_*ℓ*_*δ*_*f,ℓ*_ < 0). Because *δ*_*f,ℓ*_ ∝ *r*_*ℓ*_, this selection inflates the average per-site contribution to the bias by an amount that is proportional to the per-site contribution to the naive estimation bias. Distributing the sum and the expectation, we have

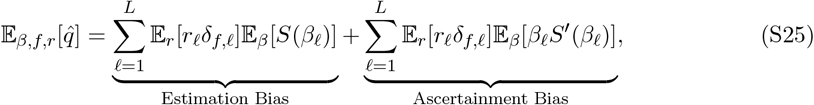

so both terms make contributions proportional to 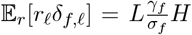. The contribution of naïve estimation bias scales with the average power,

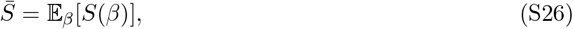

while the ascertainment bias scales with the sensitivity-weighted effect size

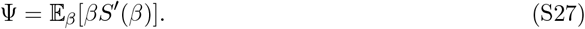

Factoring out these common terms, we obtain:

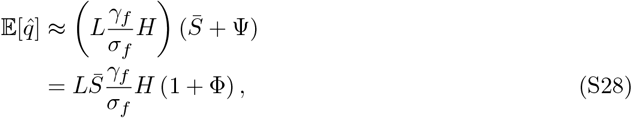

where

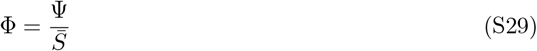

is the directional ascertainment amplification factor.

To compute the expected standardized bias, we again normalize by the standard deviation under the null. Here, we must account for the fact that the variance of *q* depends on the variance of the effect sizes of ascertained variants. We define the effect variance over ascertained sites as 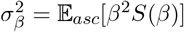. The null standard deviation is then 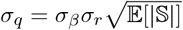.

Using the first-order approximation, 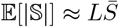, the expectation of the standardized statistic is:

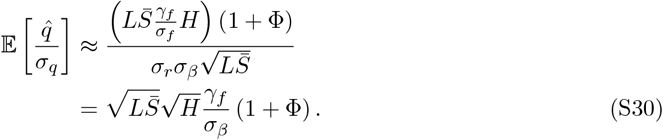

This formulation explicitly separates the effects: the numerator captures the directional bias driven by the geometric overlap in variance, *H*, and amplified by Φ. The factor Φ depends solely on the study design; for stringent thresholds, sensitivity *S*^′^ is high relative to average power 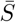, leading to a large Φ and amplified bias.

#### S1.2.3 Interaction with background stratification

The derivation above considers only the linear interaction between the directional bias *δ*_*f*_ and the ascertainment process. However, ascertainment is also affected by the total stratification load, including both the focal axis *δ*_*f*_ and any background stratification *δ*_⊥*f*_ orthogonal to the test vector.

To capture the interaction between the focal signal and this background variation, we approximate the perturbed ascertainment probability using a second-order Taylor expansion around *δ*_*f*_ = 0 and *δ*_⊥*f*_ = 0:

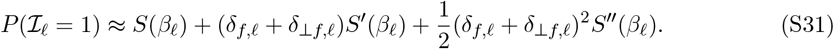

First, we consider the impact on the expected number of included SNPs, which is given by the baseline count plus a quadratic inflation term. Averaging over the distribution of effects *β*, we obtain the baseline average curvature of the power function 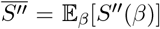, and because *δ*_*f*_ and *δ*_⊥*f*_ are uncorrelated with one another, 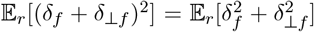, The expected SNP count can then be written as

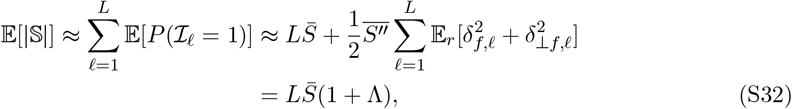

where

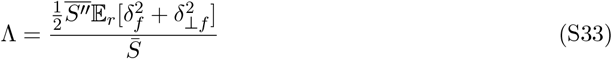

gives the average change of the ascertainment probability due to the increase in overall estimated effect variance due to stratification.

Second, we calculate the directional bias in the covariance estimator 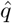. We substitute the second-order probability expansion and the expected biased effect estimate 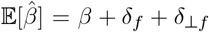 into the expectation. As before, we assume the bias *δ*_*f,ℓ*_ is independent of the true effects *β*_*ℓ*_. Averaging over the sampling noise and the distribution of causal effects *β*, we retain only the terms that correlate with the test vector *r* (i.e., terms containing *δ*_*f*_). The bias includes the standard first-order terms plus a new interaction term arising from the product of the directional estimation bias (*δ*_*f*_) and the quadratic probability inflation:

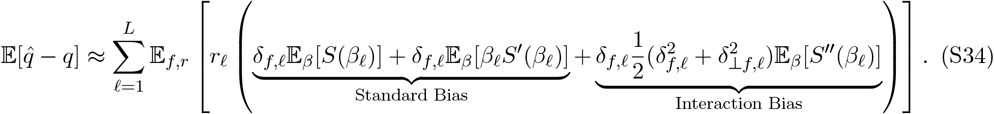

Evaluating the expectations over *β* introduces the terms 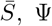 and 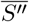 that we defined above. Distributing the expectation and the sum reveals the structure of the bias:

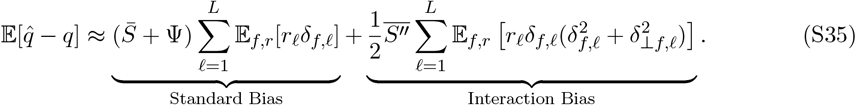

To understand the interaction bias, we separate the term into components aligned with and orthogonal to the focal axis.

For the background component, we assume that the magnitude of background stratification 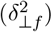 is uncorrelated with the directional signal along the target axis (*rδ*_*f*_). This allows us to factor the expectation and drop the locus-specific index:

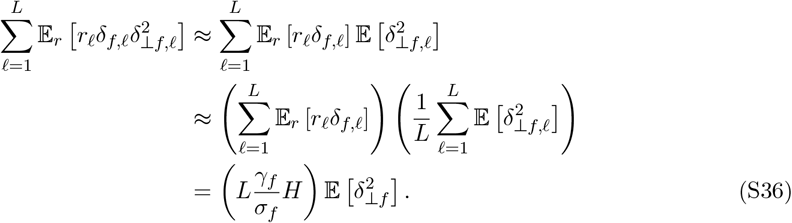

For the focal component, the term involves 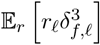. Since *δ*_*f,ℓ*_ is proportional to *r*_*ℓ*_, this term depends on the fourth moment 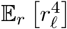. To relate this to the standard bias (which depends on the second moment 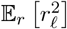), we appeal to the normal approximation for genetic drift. If *r*_*ℓ*_ follows a normal distribution, the fourth moment is related to the squared second moment by a factor of 3 (i.e., 𝔼[*r*^4^] = 3(𝔼[*r*^2^])^2^). Under this assumption, the focal interaction term is:

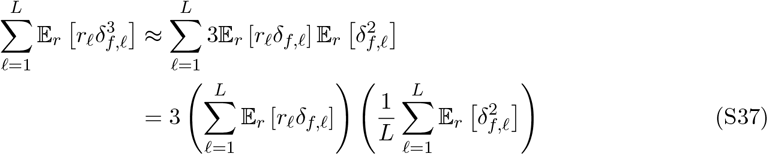

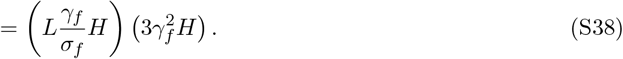

Putting these two approximations together, we can write the total bias as:

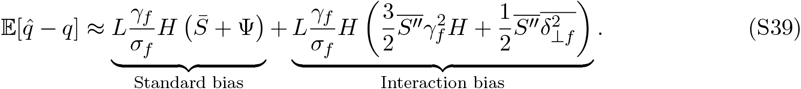

Thus, the interaction bias amplifies the naive estimation bias through a “focal” channel that scales with *H*, and a “background” channel that depends on the strength of stratification along other axes. The factor of 3 here accounts for the fact that under the normal approximation for genetic drift, some loci will have larger values of *r*^2^ (and thus larger squared bias 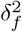); these sites will be more likely to be ascertained on the basis of their stratification bias effects, amplifying their contribution beyond what would be expected from the variance alone. No such kurtosis effect exists for the background interaction bias, as the background axes are independent of the test vector.

While the focal component is more strongly amplified due to kurtosis, in practice we expect that the total variance of stratification is dominated by background structure 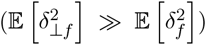, Consequently, the interaction bias is driven primarily by the orthogonal inflation of the SNP count. We therefore neglect the factor of 3 from the Gaussian approximation and write:

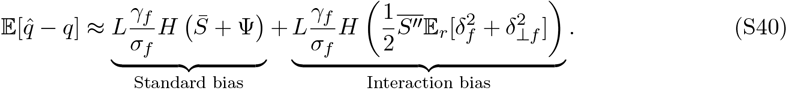

Then, recalling the definitions 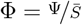 and 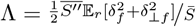 from above the total bias simplifies to:

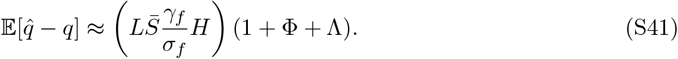

Finally, we calculate the expected standardized bias. We again define the variance of effect sizes among ascertained variants as 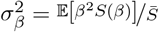. We normalize by the standard deviation under the null, 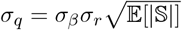. Substituting the result 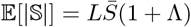:

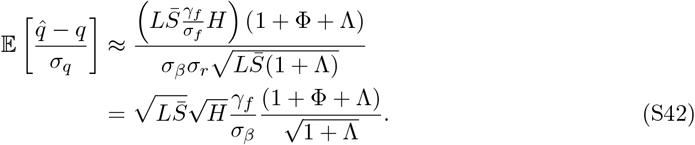

Thus, while high levels of total stratification (large Λ) inflate the noise of the PGS (the denominator,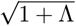), they inflate the directional bias even more strongly (the numerator, 1 + Λ), resulting in a net amplification of the standardized signal.

#### S1.2.4 Understanding the ascertainment amplification factors

The ascertainment amplification factor Φ and the background interaction factor Λ depend on the shape of the baseline ascertainment probability function *S*(*β*) and its derivatives. Here, we define this function explicitly and explore its behavior across different GWAS significance thresholds.

We assume that the test statistic for a variant with true effect *β* follows a non-central *χ*^2^ distribution with 1 degree of freedom and non-centrality parameter *λ* = *Nβ*^2^. The baseline ascertainment probability (statistical power) is the probability that this statistic exceeds the critical value *c*_*α*_ corresponding to the chosen p-value threshold *α*:

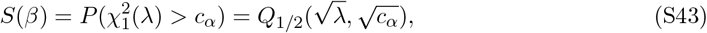

where *Q*_1/2_ is the Generalized Marcum Q-function with index *m* = 1/2.

##### Directional amplification, Φ

To understand the magnitude of the ascertainment amplification factor Φ, it is helpful to reframe it in terms of the distribution of effects among included variants. Recall that 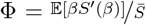, where the expectations are taken over the prior distribution of causal effects, *p*(*β*). Let 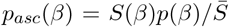 be the distribution of causal effects among variants that are successfully ascertained. We can rewrite Φ as an expectation over this ascertained distribution,

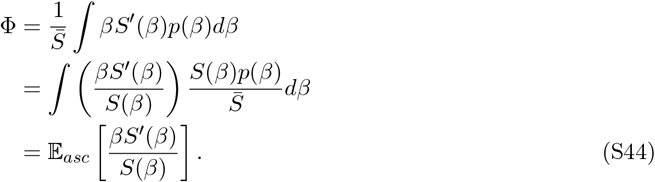

This result provides a key intuition: Φ is the average elasticity of the ascertainment probability with respect to effect size, calculated specifically among the variants included in the PGS. Formally, the quantity ^*βS*′(*β*)^*/S*(*β*) corresponds exactly to the definition of elasticity used in economics: the ratio of the percentage change in a function to the percentage change in its variable. We can see this by writing the elasticity of *S* with respect to *β* as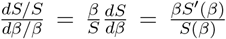. Intuitively, this metric answers the question: “If a variant’s effect size were 1% larger, by what percentage would its probability of inclusion in the score increase?” For high-power variants (*S* ≈ 1), the discovery process is inelastic; a marginal increase in effect size yields a negligible relative increase in the already-high probability of ascertainment. However, for variants on the margin of significance, ascertainment is highly elastic; a small boost in the apparent effect size (e.g., due to stratification bias) results in a disproportionately large relative increase in the probability of inclusion, thereby driving the amplification effect Φ.

Importantly, because elasticity is defined as a ratio of percentage changes, the quantity Φ is invariant to the sample size *N* of the study. While increasing sample size shifts the power curve to include variants with smaller absolute effect sizes, the functional relationship between statistical power and relative sensitivity remains constant. A variant with 50% power in a study of *N* = 10, 000 has the exact same elasticity as a variant with 50% power in a study of *N* = 500, 000, provided the significance threshold remains the same. This means that the directional amplification factor Φ is an intrinsic property of the significance threshold and the power profile of the specific ascertained variants, rather than a function of the study’s overall statistical power.

A notable feature of this relationship (visible in Figure 1d) is that for a fixed level of statistical power, the elasticity increases with the stringency of the significance threshold. We illustrate the mechanism driving this sensitivity in Figure S1. To achieve a specific level of power (e.g., *S*(*β*) = 0.5) under a stringent threshold, a variant must possess a much larger causal effect size (*β*) than is required under a lenient threshold (Figure S1a). While the local sensitivity of the power function (*S*^′^(*β*)) is determined solely by the power level itself (and is thus invariant to the threshold), the ascertainment bias depends on the product of this sensitivity and the effect size (*βS*^′^(*β*)). As shown in Figure S1b, the large causal effects required for discovery at stringent thresholds act as a multiplier on the sensitivity, increasing this product term. Consequently, for variants discovered at more stringent thresholds, the systematic nudge from stratification bias results in a disproportionately large relative increase in their ascertainment probability compared to variants found at lenient thresholds.

##### Background amplification, Λ

We can apply a similar logic to the background interaction factor, Λ. Recall that Λ captures the inflation in the number of ascertained variants due to the total stratification load, and is defined as 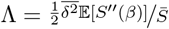, where the expectation is again over the prior distribution of causal effect (Eq. S33). Like Φ above, it can also be rewritten as an expectation over the ascertained distribution,

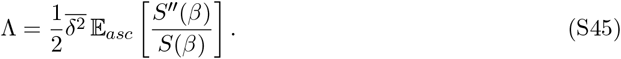

Here, the quantity ^*S*′′ (*β*)^*/S*(*β*) represents the “normalized curvature” of the ascertainment function. As shown in the bottom right panel of Figure 1, this function captures the geometric response of power to additional variance in the effect estimates. For variants with low power, the normalized curvature is positive, meaning that stratification variance pushes more variants across the threshold than it removes, resulting in a net inflation of the SNP count (Λ *>* 0). Conversely, for variants with very high power, the normalized curvature is negative, reflecting a saturation effect where added noise is more likely to push a variant below the threshold than pull a new one above it. Because polygenic scores are dominated by variants in the low-power regime, the average normalized curvature is positive, driving the characteristic inflation of test statistics associated with stratification.

### S1.3 Bias after statistical control via PCA

In Section 2.5, we introduced the use of PCA to control for stratification. Here, we derive the expected standardized bias of the corrected estimator and explicitly define the residual confounding parameter 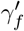.

When the top principal components are included as covariates in the GWAS, the estimated effect sizes are effectively projected onto the subspace orthogonal to these PCs. Let **P** = **I** − **UU**^⊤^ be the projection matrix onto the orthogonal complement of the top principal components. We model the corrected effect size estimate as:

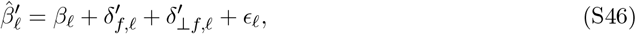

where the residual bias terms are defined by the projection of the genotypes and the environmental confounders onto this subspace.

The residual bias at locus *ℓ* arises from the covariance between the residual genotype 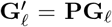 and the residual environmental confounder *c*^′^ = **P***c*. Recall that the original confounder is modeled as 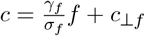. The residual confounder aligned with the target axis is therefore:

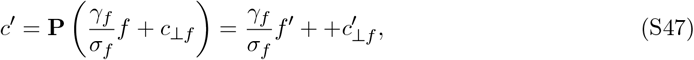

where *f*^′^ = **P***f* is the residual target axis. The systematic bias 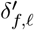 is then given by

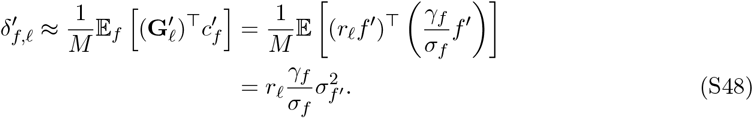

To maintain consistency with the uncorrected bias formulation (which scales with 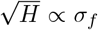), we express this residual bias in terms of the residual standard deviation in ancestry variation *σ*_*f*_′ . We define the residual confounding magnitude 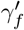 such that 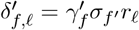, paralleling the definition of *δ*_*f,ℓ*_ in equation (S16). Equating the terms, we have

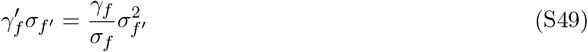

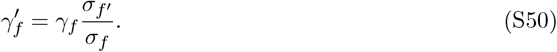

The implication of equation (S50) depends on our interpretation of the equation for the confounder, 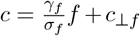 (main text equation (6)). If we interpret this as a *causal* partition of the confounder, that is if we believe that an individual’s position on the standardized ancestry axis *f* causes a linear shift in their environmental contribution (or is perfectly correlated with something that does), then equation (S50) is a “law” that relates the magnitude of the association between phenotype and the residual axis, 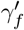, to the magnitude of the association with the original axis, *γ*_*f*_ . Under this interpretation, the values of 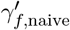 and 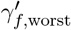 that we compute for the significant PGS-ancestry associations in the main text can be multiplied by 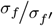 to obtain *γ*_*f*,naive_ and *γ*_*f*,worst_, i.e. a bracketing of the magnitude of the effect on the original, unresidualized axis, *f* .

However, there is no reason to think that *f* is causal. Rather, as we state in the main text, we view equation (6) is a purely statistical partition. Thus, if the environmental confounding has local structure specifically aligned with the residual axis *f*^′^, then 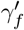 could be larger than implied by equation (S50). For this reason, it is safer to interpret 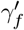 as a free parameter, and to redefine the residual confounding effect as

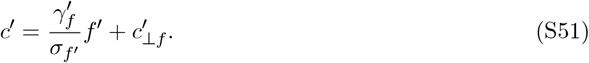

Thus, 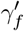 simply represents the magnitude of environmental confounding aligned with one standard deviation of the *residual* ancestry axis, and the bias is given by 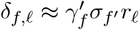.

The derivation of the bias in the PGS association test then follows the exact same logic as the uncorrected case (Section S1.2), substituting the residual stratification loads for the original loads.

First, we consider the impact on the number of included SNPs. The presence of residual stratification inflates the variance of the effect estimates, increasing the number of ascertained variants via the quadratic term in the power expansion. We define the residual stratification load parameter Λ^′^:

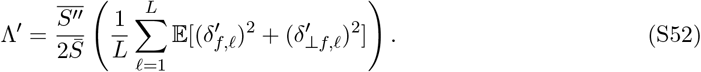

The expected number of SNPs ascertained using the corrected summary statistics is therefore:

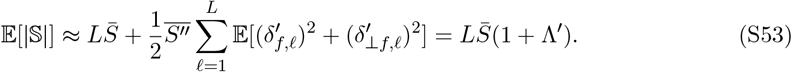

Because PCA is generally effective at removing the bulk of global genetic structure, we expect Λ^′^ ≪ Λ.

Second, we calculate the directional bias. Recognizing that 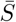 and Ψ do not depend on the stratification bias terms, we can plug the residual bias terms, 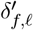 and 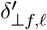 into Equation S34 to get:

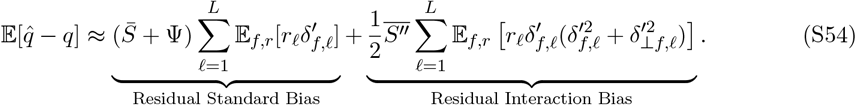

Following the same steps as above, the total bias becomes:

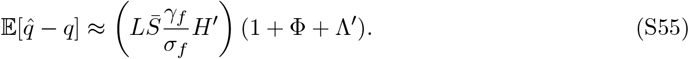

Finally, we calculate the expected standardized bias. We normalize by the standard deviation under the null, accounting for the residual inflation of the SNP count: 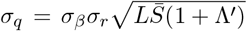. The expectation of the standardized statistic is:

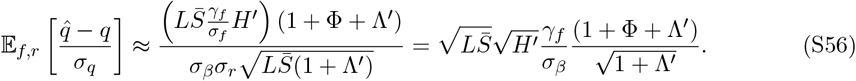

This confirms that while PCA reduces the bias by reducing the variance explained (from *H* to *H*^′^) and by reducing the background ascertainment inflation (from Λ to Λ^′^), the directional amplification due to ascertainment (Φ) continues to operate on whatever residual signal remains.

**Figure S1:**
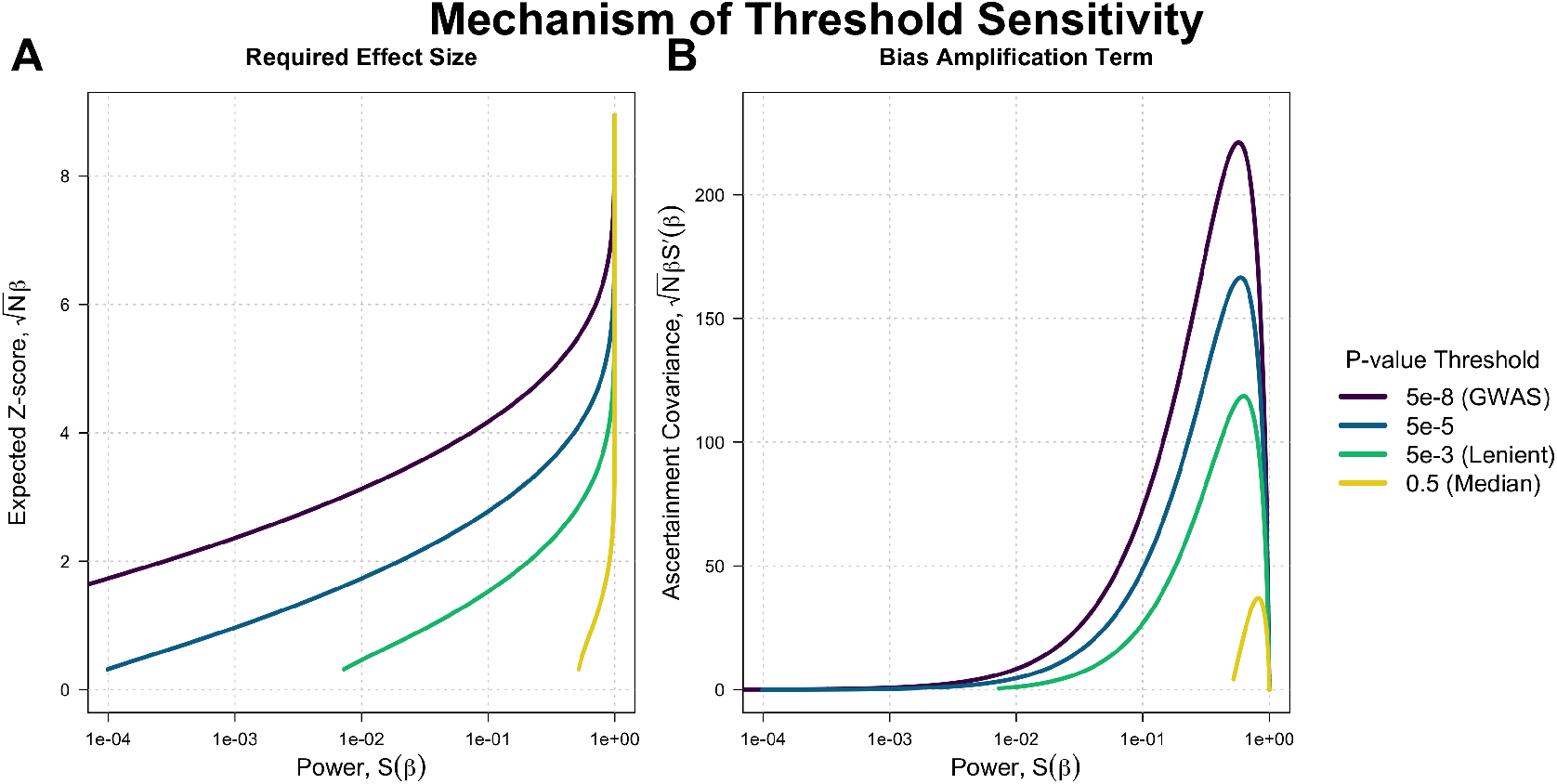
Mechanism of threshold sensitivity. We decompose the components of ascertainment elasticity to explain why stringent thresholds yield higher amplification factors (Φ). (a) The required causal effect size (*β*) required to attain a given statistical power, plotted against that power. Note that effect sizes are plotted on the expected Z-score scale 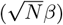 to remove the effect of sample size. To achieve the same level of power, stringent thresholds (purple) require much larger causal effect sizes than lenient thresholds (yellow/green). (b) The Z-score scaled ascertainment amplification term 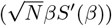 plotted against power. Because elasticity is defined as ^*βS*′(*β*)^*/S*, and the sensitivity *S*^′^ is determined solely by the power level *S* (and is thus invariant to the threshold), the larger effect sizes required by stringent thresholds (shown in a) act as a multiplier on the sensitivity. This results in the higher elasticity curves observed in Figure 1d.

**Table S1:** Sample sizes for different groups in the combined HGDP 1kGP dataset.

| HGDP1kGP Prediction Panel |  |  |
| --- | --- | --- |
| Group | Code | Sample Size |
| East Asia | eas | 825 |
| Africa | afr | 1003 |
| South Asia | sas | 790 |
| Non-Finish Europe | nfe | 689 |
| America | amr | 552 |
| Eurasia | eurasia | 2224 |
| Europe | eur | 510 |
| Sardinia | sdi | 27 |

**Figure S2:**
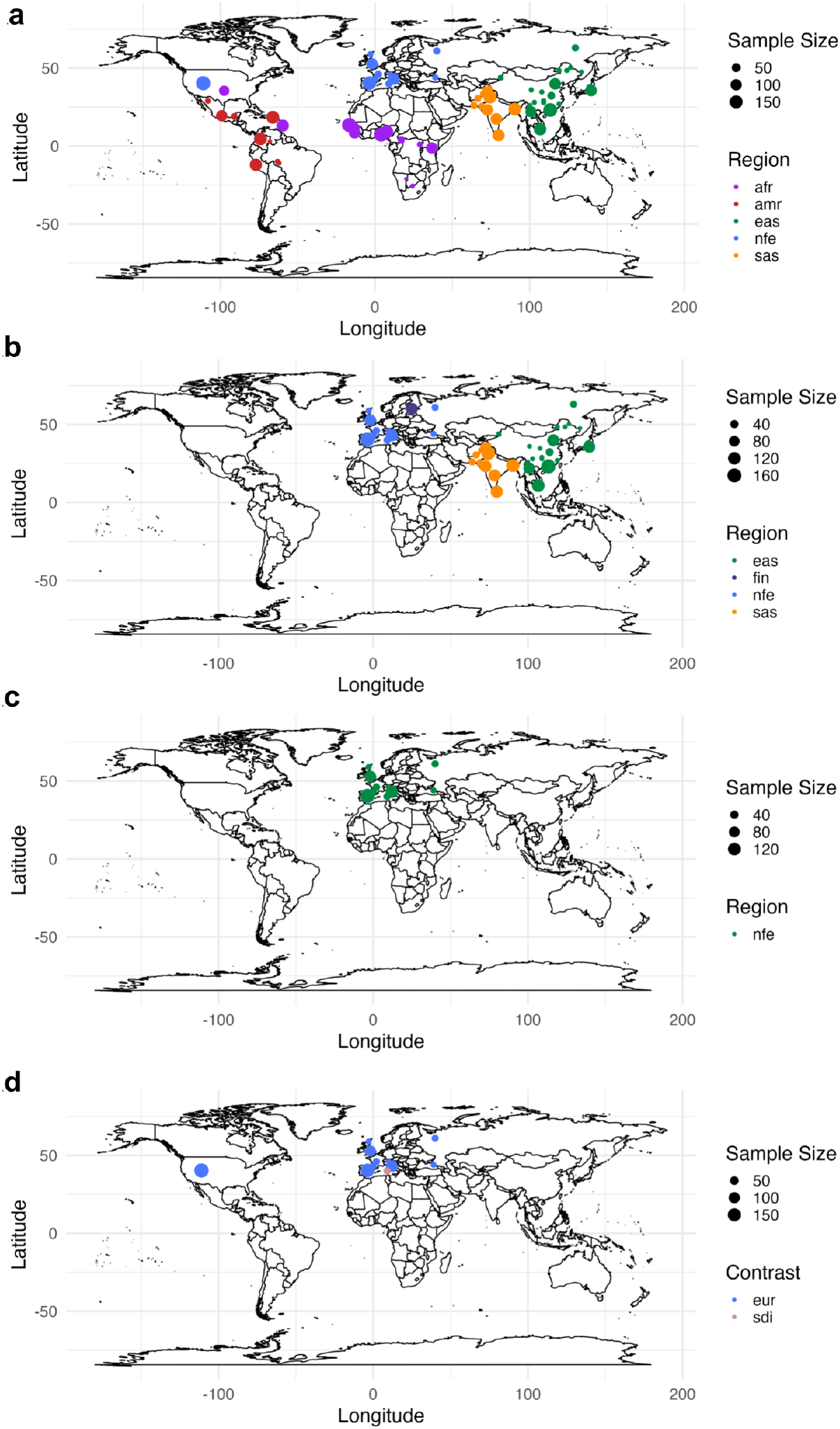
HGDP and 1kGP test panels. (a) Panel used to compute pairwise continental allele frequency differences. (b) Eurasian panel. (c) Non-Finnish European panel. (d) Sardinian panel that includes all Sardinian and non-Finnish Europeans.

**Figure S3:**
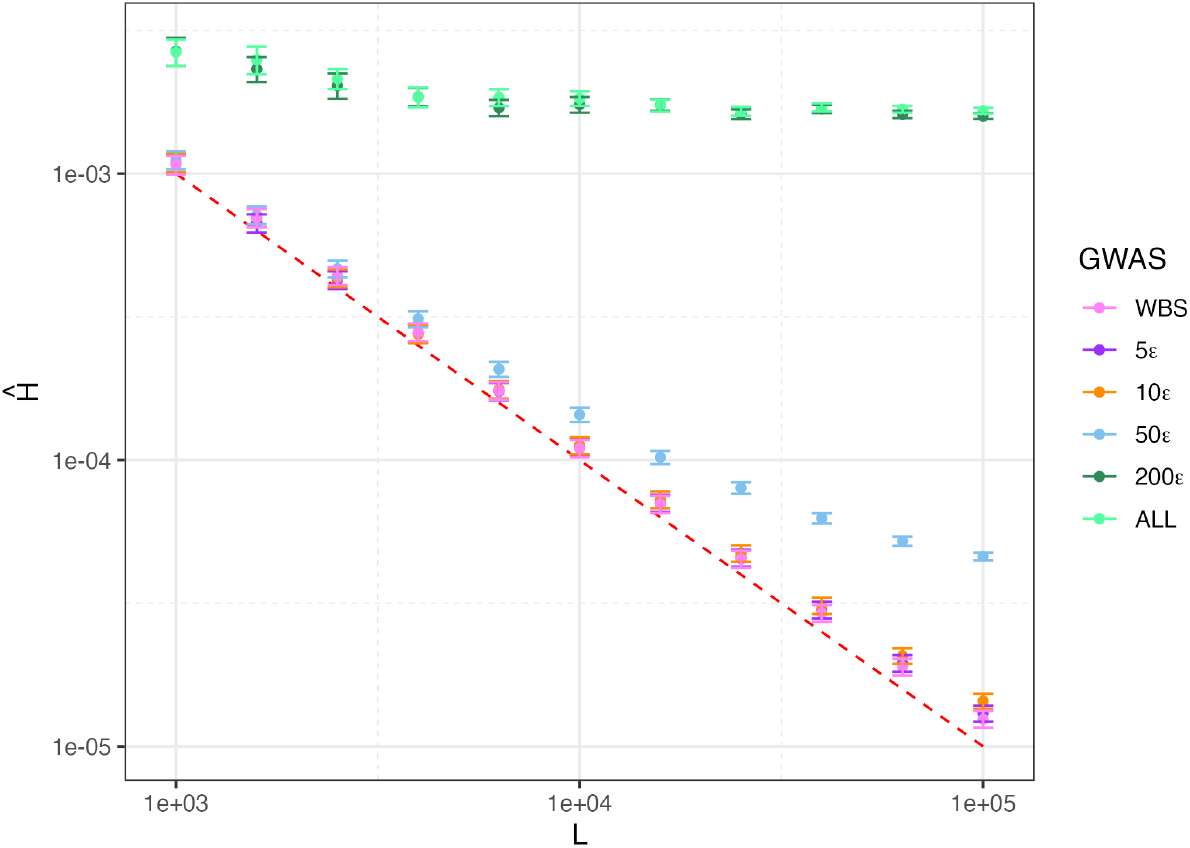
Sensitivity of uncorrected susceptibility estimates to SNP set size. We re-estimated *Ĥ* for a single representative contrast (East Asia vs. South Asia) using subsets of the UK Biobank PC markers ranging from *L* = 1, 000 to *L* = 100, 000 SNPs. Results are shown for each of the six GWAS panels. The dashed red line indicates the theoretical limit of detection due to sampling noise (≈ 1*/L*). As the number of SNPs decreases (increasing the physical distance between markers and reducing any potential LD), the estimates for the homogeneous panels (WBS, 5*ϵ*, 10*ϵ*) converge to the detection limit, but remain distinguishable from it with only *L* ≈ 2 × 10^4^ SNPs, indicating that the signal is unlikely to be the result of inflation due to LD. In the more diverse panels, the signal remains robustly detectable even with fewer markers.

**Table S2:** Sample sizes used to construct prediction panels and estimate 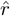 for different countries of birth within the British Isles.

| Within the British Isles Country of Birth Prediction Panel |  |  |
| --- | --- | --- |
| Country | Code | Sample Size |
| England | ENG | 142215 |
| Northern Ireland | NI | 1120 |
| Republic of Ireland | RoI | 1802 |
| Scotland | SCT | 14591 |
| Wales | WAL | 8125 |

**Figure S4:**
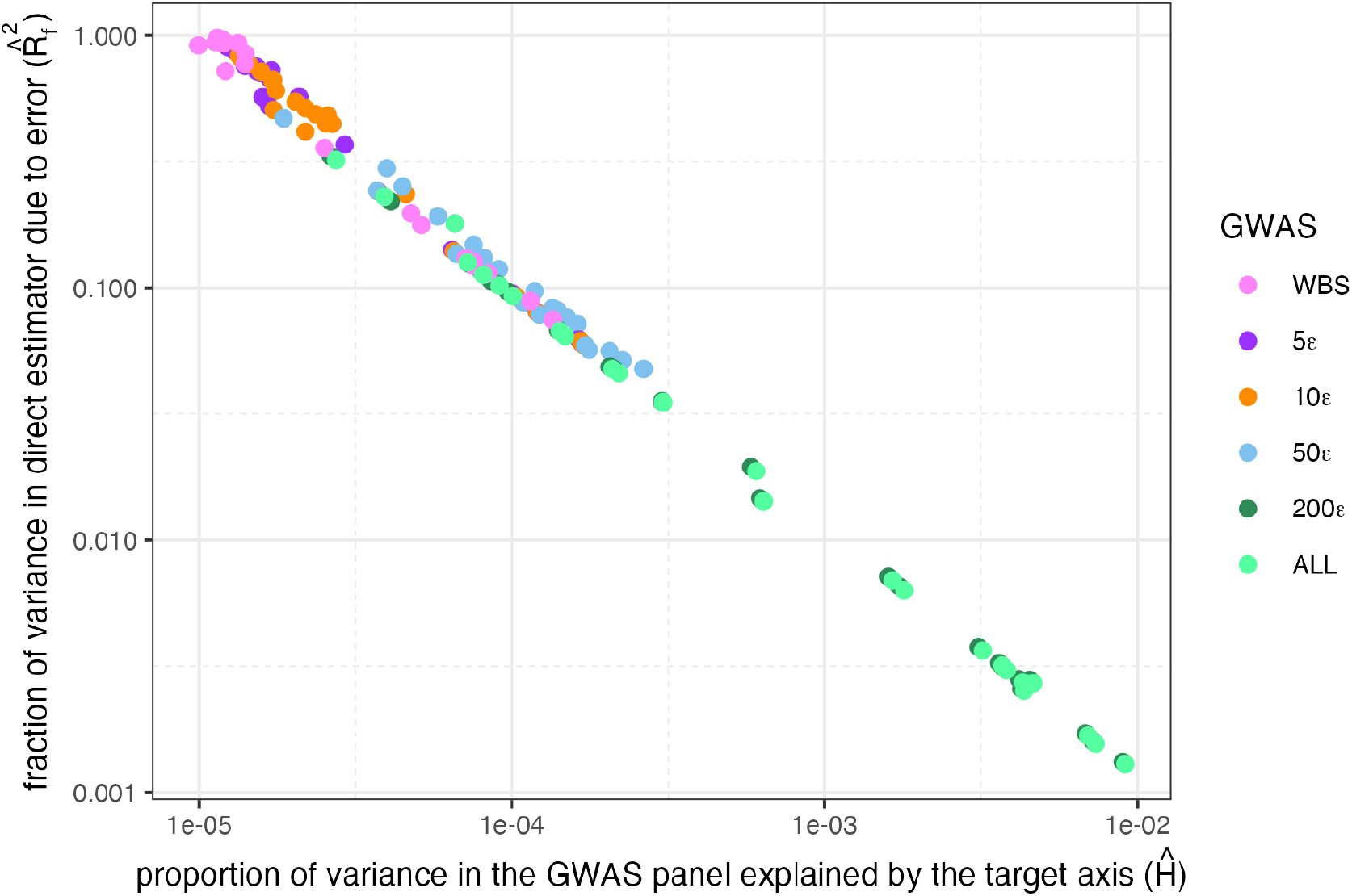
Estimation error in the target axis decreases with signal strength. We estimated the target ancestry axis 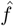 for each GWAS panel/contrast pair and used a weighted block jackknife to quantify the proportion of variance in this estimator attributable to sampling noise 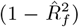, The estimation error is inversely related to the strength of the stratification signal 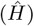, meaning that target axes which explain a larger proportion of genetic variance are estimated with higher precision.

**Figure S5:**
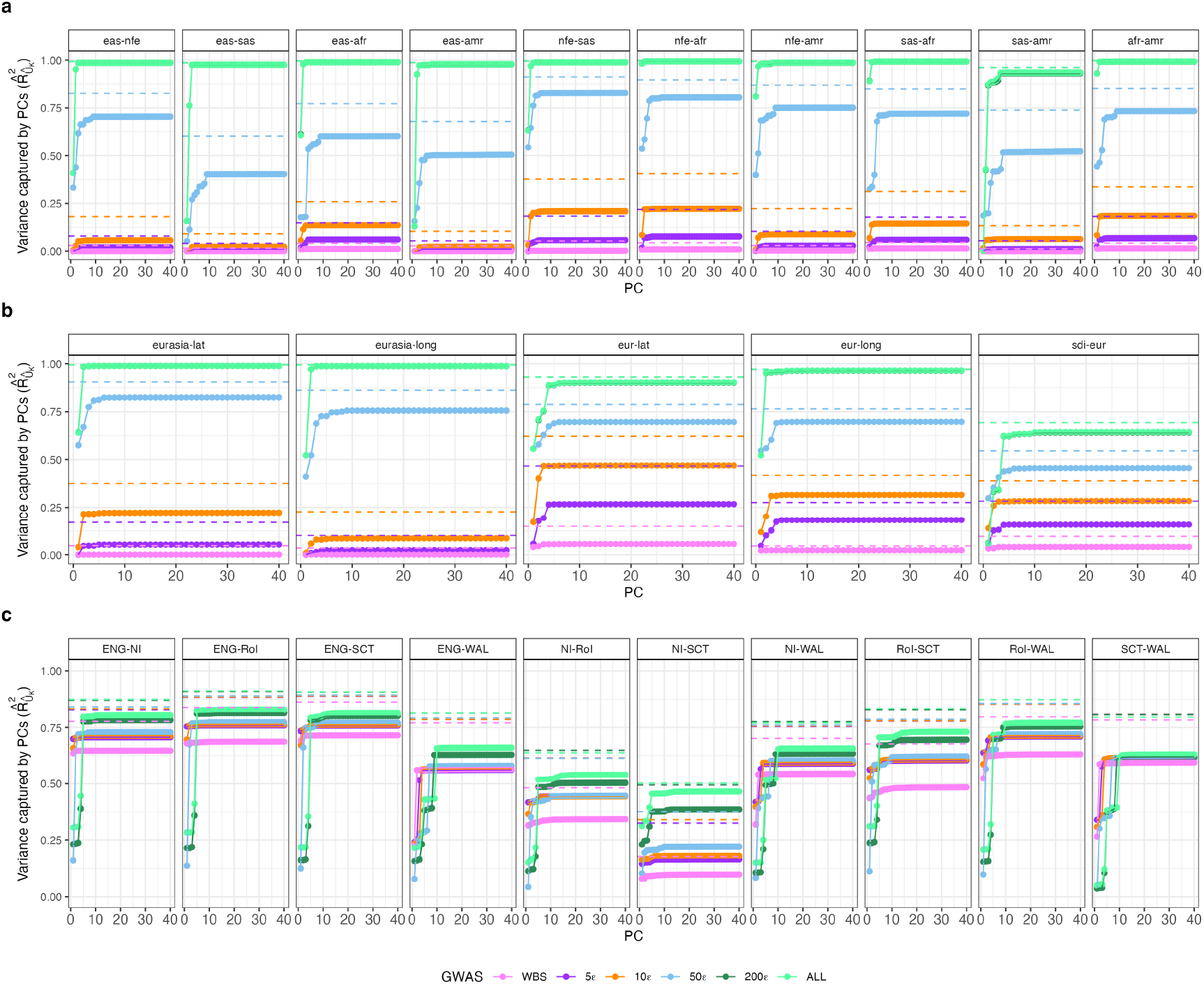
Cumulative variance in the target axis captured by common variant PCs. Each point represents 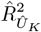, the fraction of variance in the estimated target axis 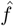 that is explained by the top *K* common variant principal components (*K* = 1 … 40) within a specific GWAS panel. The dashed horizontal line indicates 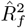, the estimated reliability of the target axis vector itself (i.e., the fraction of variance in 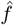 attributable to true genetic signal rather than estimation noise). Because the PCs cannot explain noise, 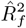 serves as the theoretical upper bound for 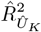.

**Figure S6:**
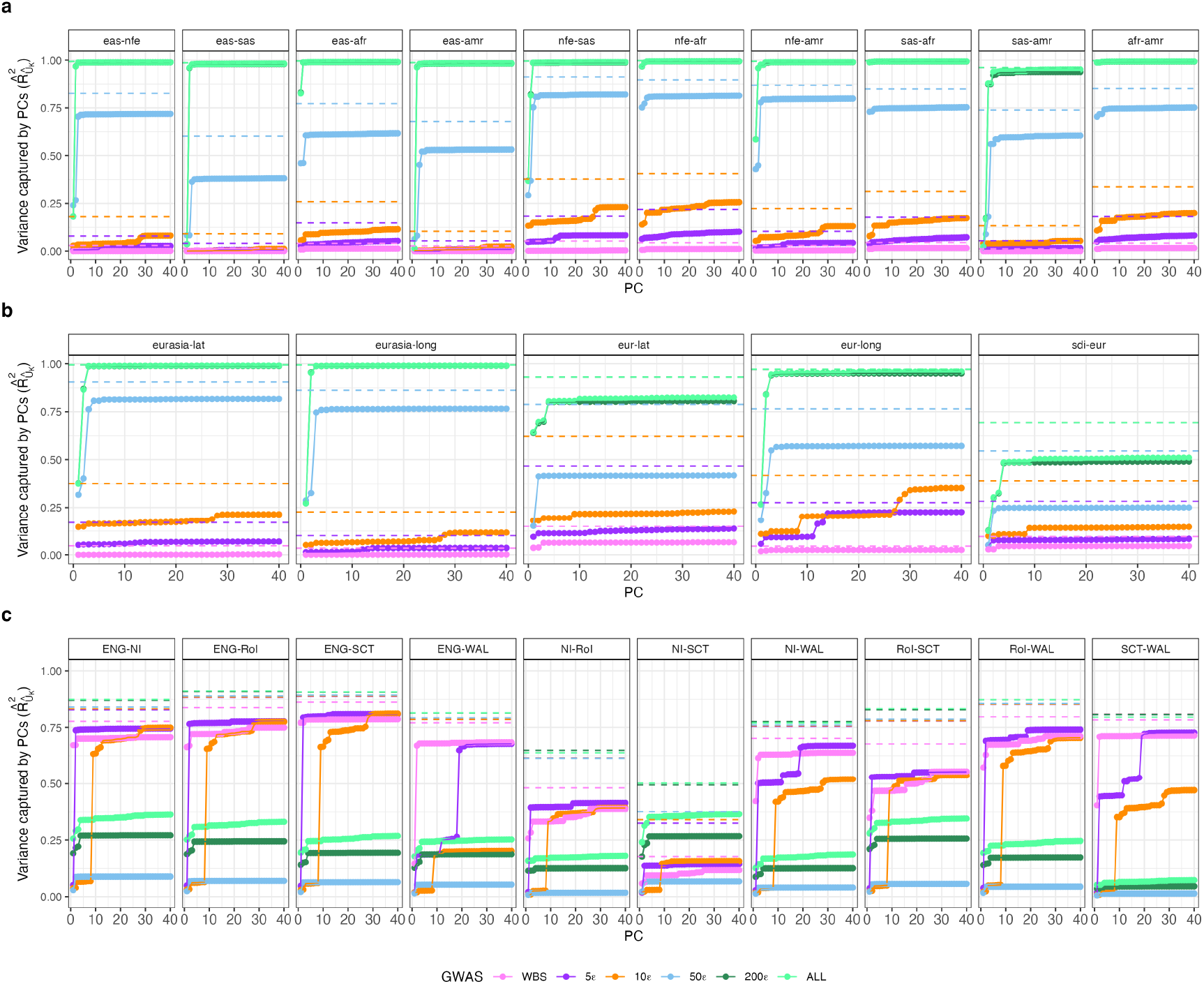
Cumulative variance in the target axis captured by rare variant PCs. Each point represents 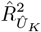, the fraction of variance in the estimated target axis 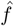 that is explained by the top *K* rare variant principal components (*K* = 1 … 40) within a specific GWAS panel. The dashed horizontal line indicates 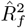, the estimated reliability of the target axis vector itself (i.e., the fraction of variance in 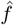 attributable to true genetic signal rather than estimation noise). Because the PCs cannot explain noise, 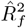 serves as the theoretical upper bound for 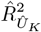.

**Figure S7:**
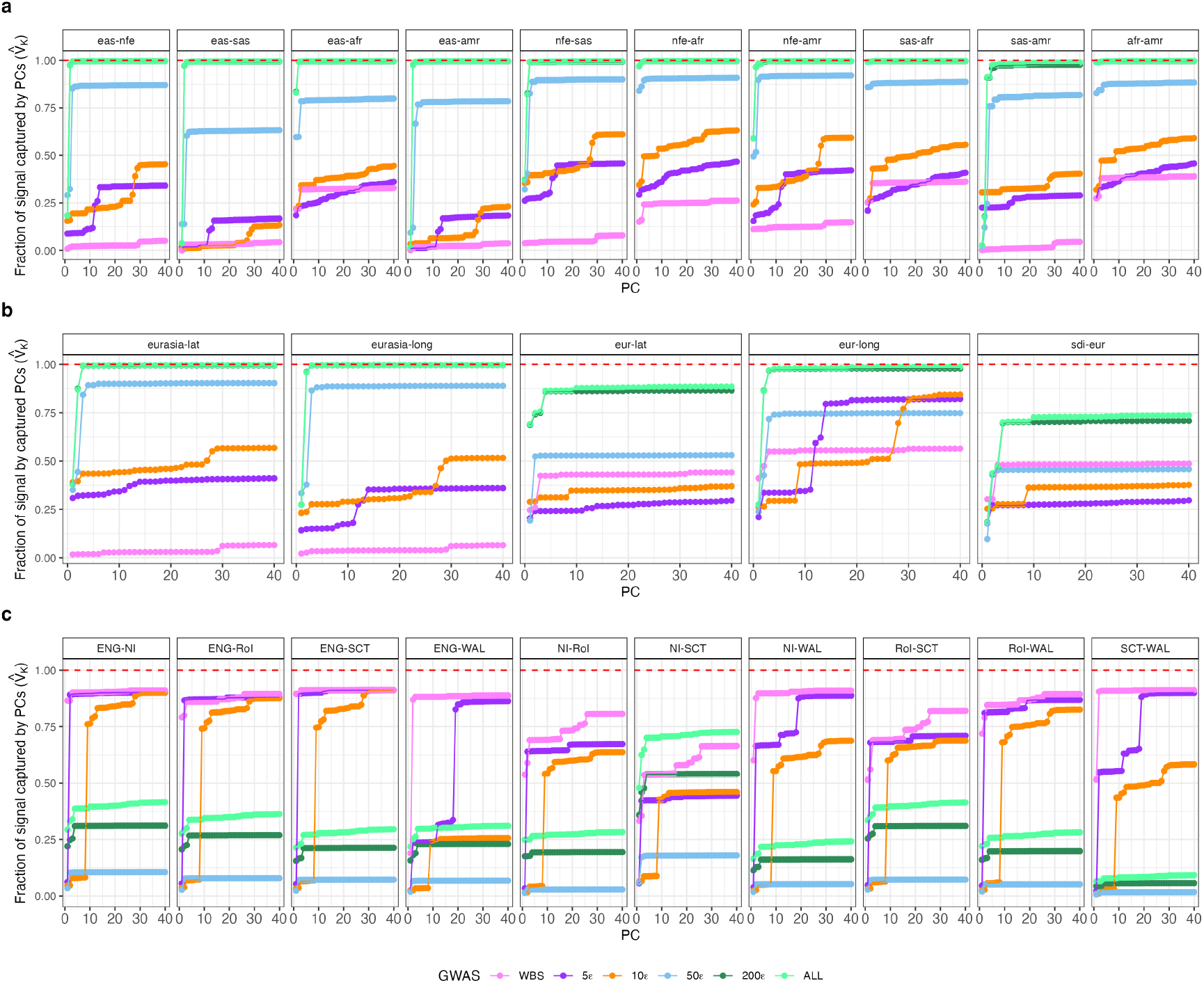
Efficacy of rare variant PCs in capturing target axes. We estimated 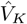 the proportion of the explainable variance in the target axis that is captured by the top *K* rare variant principal components 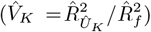. Plots show the cumulative efficacy for *K* = 1 … 40 rare variant PCs used in isolation, without the prior correction for common variant PCs that is applied in Figure 3.

**Figure S8:**
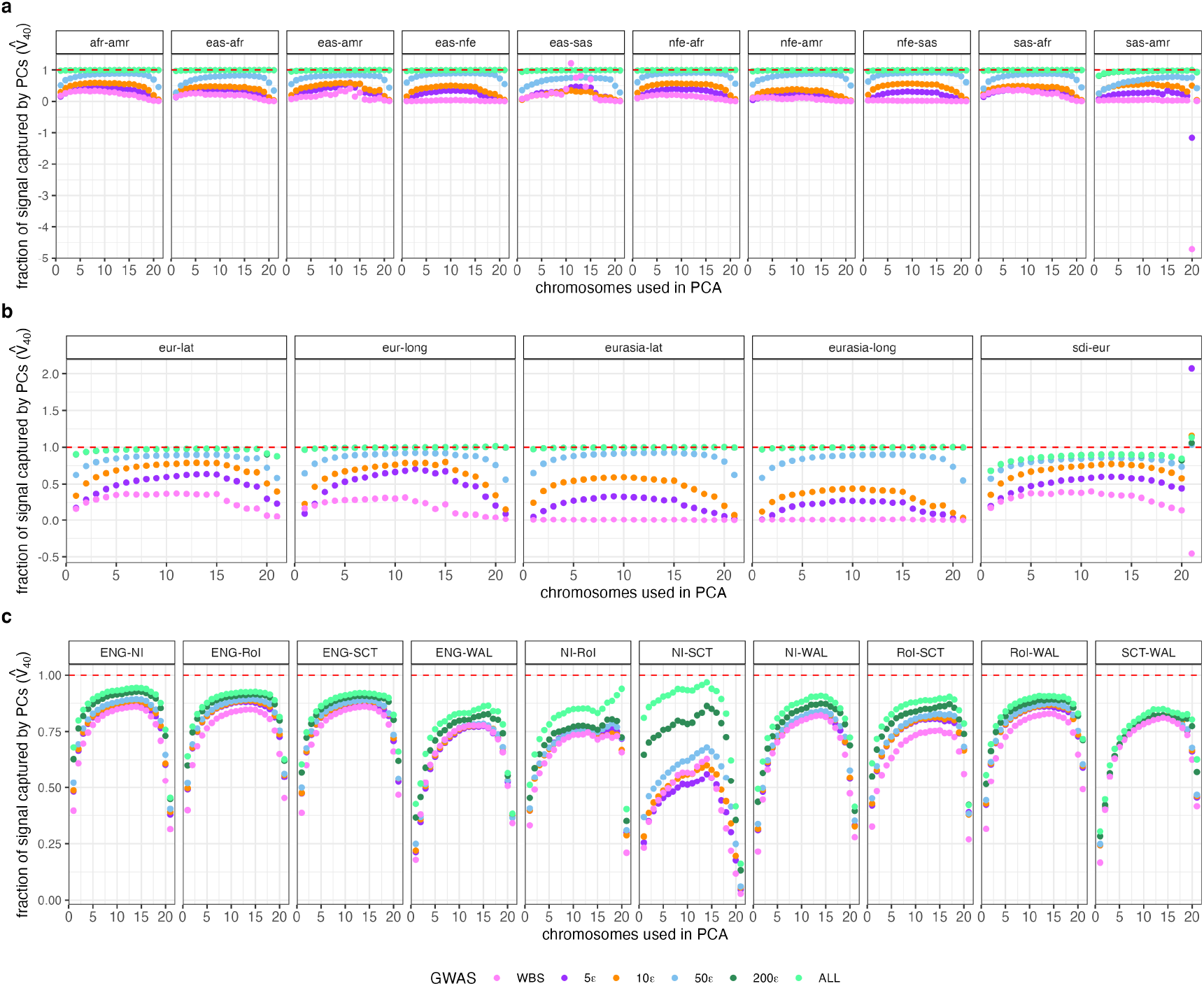
Robustness of PCA efficacy estimates to sample splitting. We calculated the PCA efficacy metric 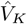 using a split-sample strategy to avoid the overfitting bias that arises when the target axis 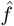 and the principal components 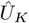 are estimated from the same genotypes. In the primary analysis, we split chromosomes into odd and even sets. Here, we vary the split proportion by using only the first *C* chromosomes (*x*-axis) to compute the PCs, while using the remaining chromosomes to estimate the target axis. The resulting 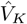 estimates are relatively stable across different splitting strategies, provided that a sufficient number of chromosomes (≳ 5) are used for each component estimation.

**Figure S9:**
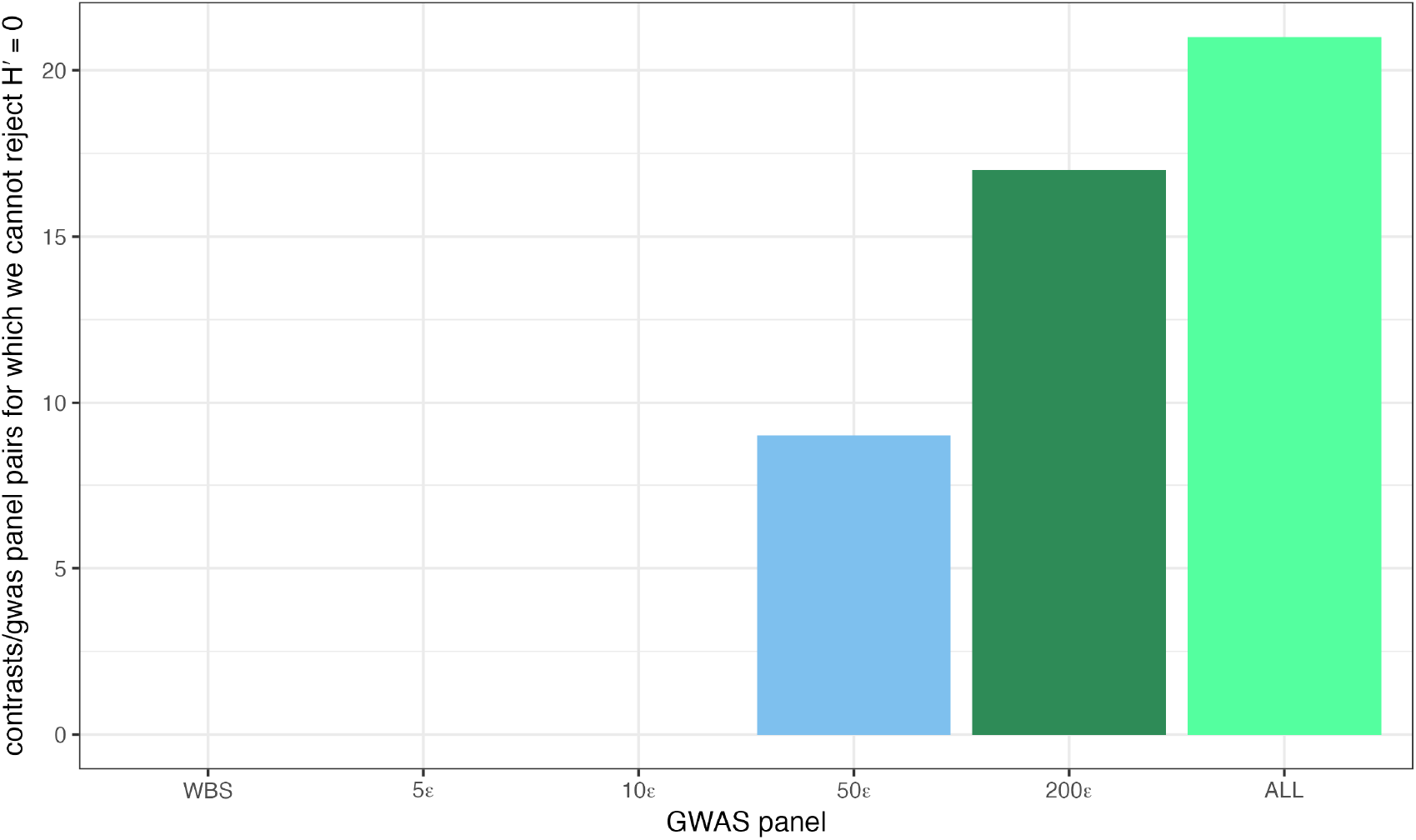
Detection of residual structure varies by study design. We tested the null hypothesis of zero residual susceptibility (*Ĥ*^′^ = 0) for each of the 25 genotype contrasts after correcting for 40 common and 40 rare principal components. Bars show the number of contrasts (out of 25) for which we failed to reject the null hypothesis (*p >* 0.05 after Bonferroni correction). The diverse ALL panel exhibits the highest number of contrasts for which residual stratification is indistinguishable from sampling noise. This suggests that the increased precision of PC estimation in large, diverse cohorts reduces the signal below the limit of detection for the majority of tested axes, whereas residual structure remains statistically detectable for all contrasts in the WBS and other narrowly sampled panels.

**Figure S10:**
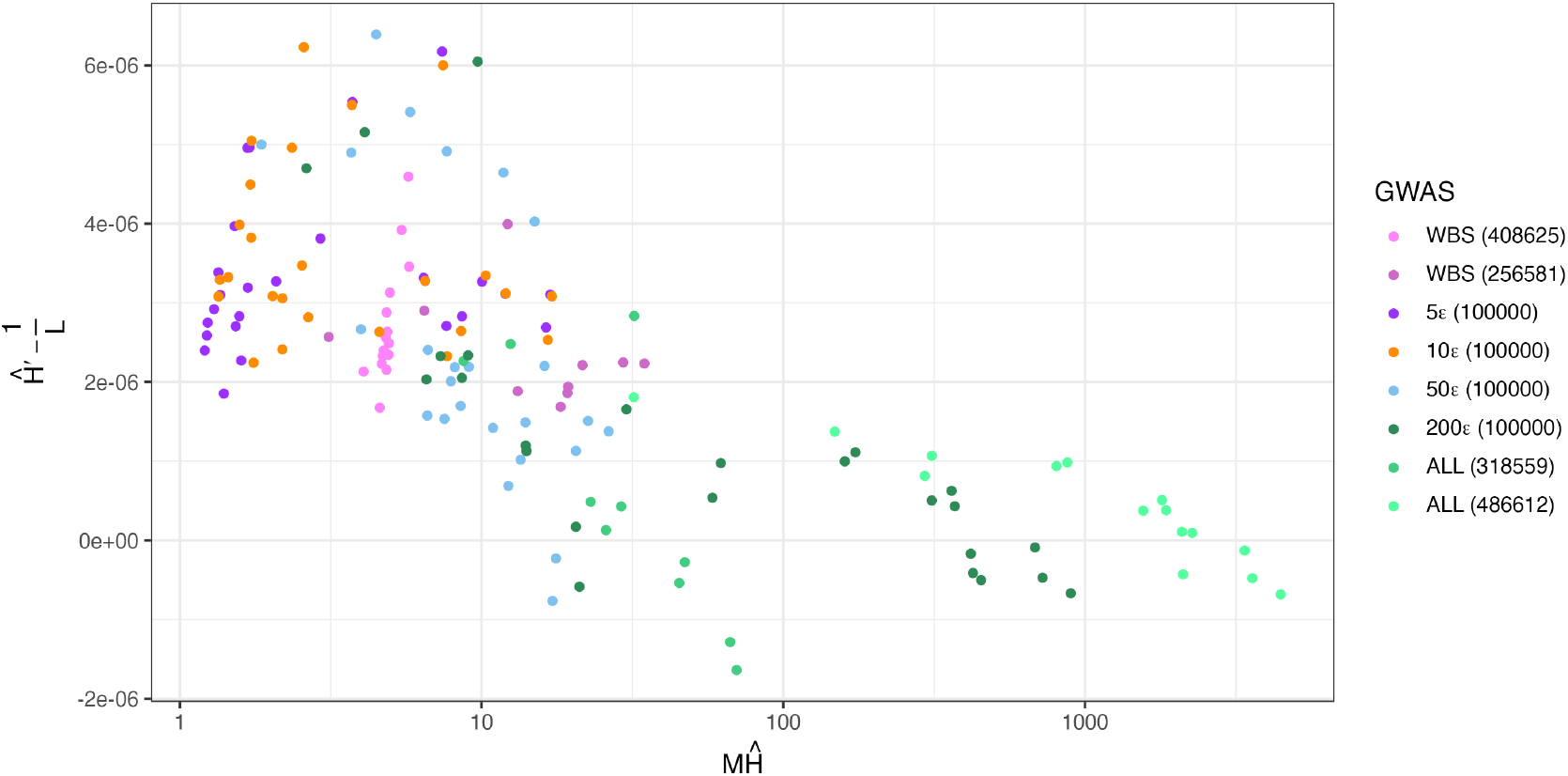
Residual susceptibility decreases with GWAS sample size. For each GWAS panel/contrast pair, we plot the estimated residual susceptibility *Ĥ*^′^ (after correcting for 40 common and 40 rare PCs) against the product of the GWAS sample size (*M*) and the initial uncorrected susceptibility (*Ĥ*). This quantity, *M Ĥ*, serves as a proxy for the total information available to estimate the target axis via PCA. As predicted by theory [11, 49**?** ], higher information content (larger *M Ĥ*) generally leads to lower residual structure. Note that British Isle country of birth contrasts have smaller effective sample sizes in the ALL and WBS panels because overlapping individuals from the prediction set were removed.

**Table S3:** UKBB phenotypes used for polygenic score association tests.

| Phenotype | Code |
| --- | --- |
| Standing Height | 50 |
| Alkaline Phosphate | 30610 |
| Aspartate Aminotransferase | 30650 |
| Basophil Percentage | 30220 |
| Body Weight | 21002 |
| Cholesterol | 30690 |
| Eosinophil Percentage | 30210 |
| Glucose | 30740 |
| HbA1c | 30750 |
| HDL | 30760 |
| LDL | 30780 |
| Mean Corpuscular Haemoglobin Concentration (MCHC) | 30060 |
| Mean Corpuscular Volume (MCV) | 30040 |
| Platelet Count | 30080 |
| Systolic Blood Pressure (SBP) | 4080 |
| Total Protein | 30860 |
| Triglycerides | 30870 |

**Figure S11:**
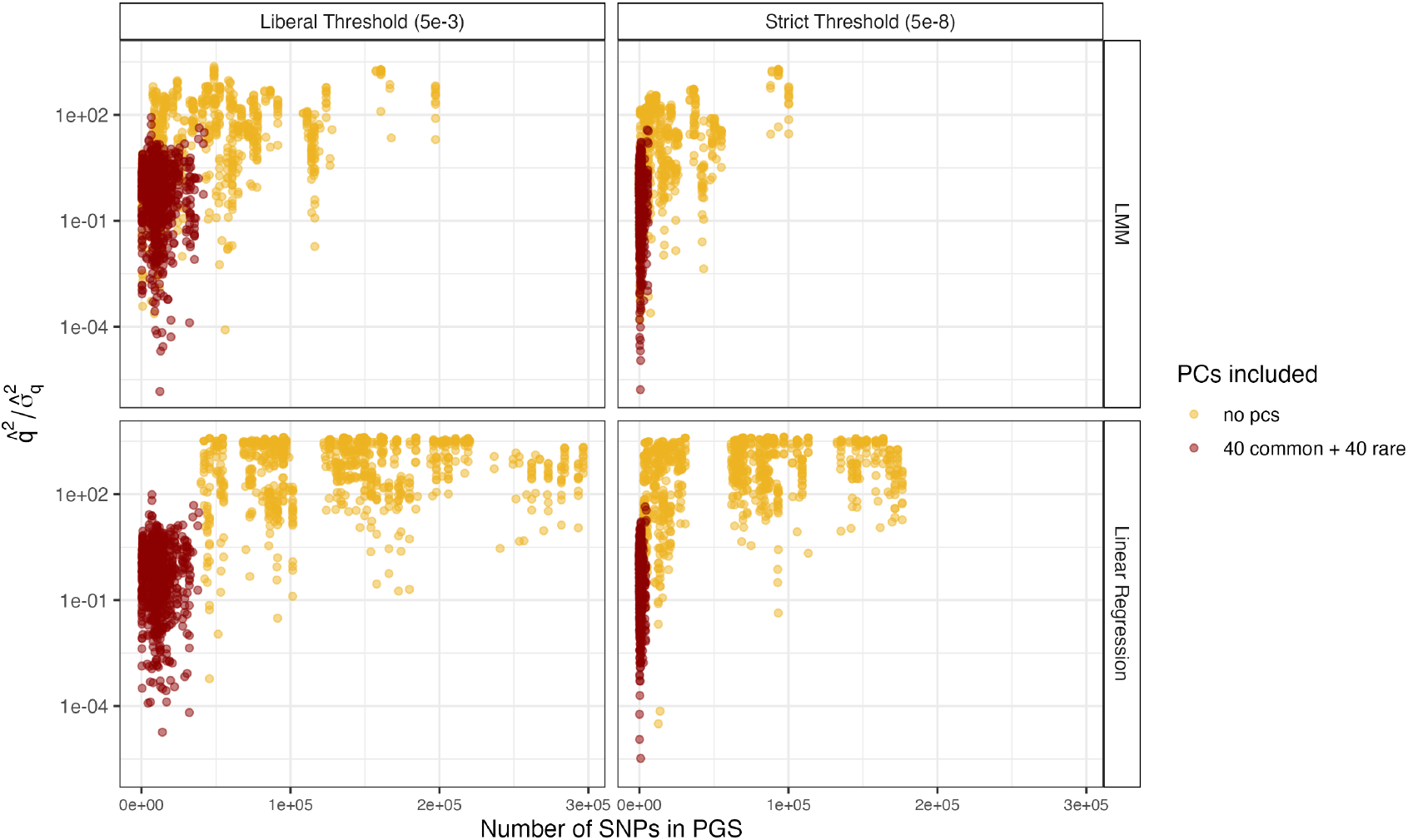
Total stratification load inflates the number of ascertained SNPs. We plot the standardized squared test statistics 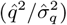 against the number of SNPs included in each polygenic score. When no PCs are included in the GWAS (yellow), the background stratification load significantly increases the variance of the effect size estimates. This inflation pushes a large number of null variants across the significance threshold, drastically increasing the number of SNPs ascertained for the score (x-axis) compared to the PC-corrected models (red). This global inflation of the SNP count corresponds to the theoretical parameter Λ.

**Table S4:** Significant 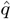 values across all methodologies.

| Dataset | GWAS | Contrast | GWAS type | Threshold | Phenotype | $\hat{H}'$ | $\hat{q}/\hat{\sigma}_q$ | p-value | S |
| --- | --- | --- | --- | --- | --- | --- | --- | --- | --- |
| HGDP1KG | ALL | eur-lat | LR | strict | Standing Height | 8.642e-06 | 4.260222 | 2.042e-05 | 4496 |
| HGDP1KG | ALL | sdi-eur | LR | strict | Standing Height | 1.083e-05 | -5.940780 | 2.837e-09 | 4519 |
| HGDP1KG | ALL | sdi-eur | LR | lenient | Standing Height | 1.083e-05 | -5.483959 | 4.159e-08 | 38428 |
| HGDP1KG | ALL | sas-amr | LMM | strict | MCV | 9.837e-06 | -4.020751 | 5.801e-05 | 1169 |
| HGDP1KG | ALL | eur-lat | LMM | strict | Standing Height | 8.642e-06 | 3.942111 | 8.077e-05 | 5900 |
| HGDP1KG | ALL | sdi-eur | LMM | strict | Standing Height | 1.083e-05 | -5.906332 | 3.498e-09 | 5930 |
| HGDP1KG | ALL | sdi-eur | LMM | lenient | Body Weight | 1.083e-05 | -3.885159 | 1.023e-04 | 28926 |
| HGDP1KG | ALL | eur-lat | LMM | lenient | Standing Height | 8.642e-06 | 3.903575 | 9.478e-05 | 41508 |
| HGDP1KG | ALL | sdi-eur | LMM | lenient | Standing Height | 1.083e-05 | -5.630671 | 1.795e-08 | 42147 |
| HGDP1KG | WBS | sdi-eur | LR | strict | Body Weight | 1.262e-05 | -4.069407 | 4.713e-05 | 1115 |
| HGDP1KG | WBS | sas-amr | LR | strict | MCV | 1.139e-05 | -3.905659 | 9.397e-05 | 937 |
| HGDP1KG | WBS | eur-lat | LR | strict | Standing Height | 1.192e-05 | 3.998529 | 6.374e-05 | 3822 |
| HGDP1KG | WBS | sdi-eur | LR | strict | Standing Height | 1.262e-05 | -6.631391 | 3.325e-11 | 3843 |
| HGDP1KG | WBS | eur-lat | LR | lenient | Standing Height | 1.192e-05 | 4.759427 | 1.941e-06 | 34417 |
| HGDP1KG | WBS | sdi-eur | LR | lenient | Standing Height | 1.262e-05 | -6.946027 | 3.757e-12 | 34988 |
| HGDP1KG | WBS | sas-amr | LMM | strict | MCV | 1.139e-05 | -4.109398 | 3.967e-05 | 1021 |
| HGDP1KG | WBS | eur-lat | LMM | strict | Standing Height | 1.192e-05 | 4.182623 | 2.882e-05 | 5119 |
| HGDP1KG | WBS | sdi-eur | LMM | strict | Standing Height | 1.262e-05 | -6.142990 | 8.098e-10 | 5137 |
| HGDP1KG | WBS | eur-lat | LMM | lenient | Standing Height | 1.192e-05 | 4.543296 | 5.538e-06 | 38153 |
| HGDP1KG | WBS | sdi-eur | LMM | lenient | Standing Height | 1.262e-05 | -6.513829 | 7.326e-11 | 38717 |
| InUKBB | WBS | ENG-SCT | LR | lenient | SBP | 9.128e-06 | 5.144775 | 2.678e-07 | 5170 |
| InUKBB | WBS | ENG-SCT | LMM | lenient | SBP | 9.128e-06 | 4.397962 | 1.093e-05 | 4938 |
| InUKBB | WBS | RoI-SCT | LMM | lenient | SBP | 1.089e-05 | 4.232598 | 2.310e-05 | 4938 |
| InUKBB | ALL | ENG-NI | LR | lenient | SBP | 6.615e-06 | 4.914621 | 8.895e-07 | 6985 |
| InUKBB | ALL | ENG-SCT | LR | lenient | SBP | 5.607e-06 | 9.870656 | 5.580e-23 | 6985 |
| InUKBB | ALL | NI-WAL | LR | lenient | SBP | 7.375e-06 | -4.345840 | 1.387e-05 | 6985 |
| InUKBB | ALL | RoI-SCT | LR | lenient | SBP | 9.724e-06 | 4.945858 | 7.581e-07 | 6985 |
| InUKBB | ALL | SCT-WAL | LR | lenient | SBP | 7.019e-06 | -8.146087 | 3.759e-16 | 6985 |
| InUKBB | ALL | ENG-SCT | LMM | lenient | SBP | 5.607e-06 | 9.221873 | 2.920e-20 | 6620 |
| InUKBB | ALL | RoI-SCT | LMM | lenient | SBP | 9.724e-06 | 5.147751 | 2.636e-07 | 6620 |
| InUKBB | ALL | SCT-WAL | LMM | lenient | SBP | 7.019e-06 | -7.289962 | 3.100e-13 | 6620 |

## References

1. Sohail M, Maier RM, Ganna A, Bloemendal A, Martin AR, Turchin MC, et al. Polygenic adaptation on height is overestimated due to uncorrected stratification in genome-wide association studies. Elife. 2019;8.

2. Berg JJ, Harpak A, Sinnott-Armstrong N, Joergensen AM, Mostafavi H, Field Y, et al. Reduced signal for polygenic adaptation of height in UK Biobank. Elife. 2019;8.

3. Blanc J, Berg JJ. Testing for differences in polygenic scores in the presence of confounding. Genetics. 2025;230(2).

4. Ding Y, Hou K, Xu Z, Pimplaskar A, Petter E, Boulier K, et al. Polygenic scoring accuracy varies across the genetic ancestry continuum. Nature. 2023;618(7966):774–781.

5. International Schizophrenia Consortium, Purcell SM, Wray NR, Stone JL, Visscher PM, O’Donovan MC, et al. Common polygenic variation contributes to risk of schizophrenia and bipolar disorder. Nature. 2009;460(7256):748–752.

6. Price AL, Patterson NJ, Plenge RM, Weinblatt ME, Shadick NA, Reich D. Principal components analysis corrects for stratification in genome-wide association studies. Nat Genet. 2006;38(8):904–909.

7. Kang HM, Sul JH, Service SK, Zaitlen NA, Kong SY, Freimer NB, et al. Variance component model to account for sample structure in genome-wide association studies. Nat Genet. 2010;42(4):348–354.

8. Loh PR, Tucker G, Bulik-Sullivan BK, Vilhjálmsson BJ, Finucane HK, Salem RM, et al. Efficient Bayesian mixed-model analysis increases association power in large cohorts. Nat Genet. 2015;47(3):284–290.

9. Jiang L, Zheng Z, Qi T, Kemper KE, Wray NR, Visscher PM, et al. A resource-efficient tool for mixed model association analysis of large-scale data. Nat Genet. 2019;51(12):1749–1755.

10. Zhou W, Nielsen JB, Fritsche LG, Dey R, Gabrielsen ME, Wolford BN, et al. Efficiently controlling for case-control imbalance and sample relatedness in large-scale genetic association studies. Nat Genet. 2018;50(9):1335–1341.

11. Bloemendal A, Chen C. PCA and stratification in GWAS / A primer on random matrix theory; 2019. https://www.youtube.com/watch?v=B7ub92OLw1g.

12. Hoffman GE. Correction: Correcting for population structure and kinship using the linear mixed model: Theory and extensions. PLoS One. 2013;8(12).

13. Zhang Y, Pan W. Principal component regression and linear mixed model in association analysis of structured samples: competitors or complements? Genet Epidemiol. 2015;39(3):149–155.

14. Schraiber JG, Edge MD, Pennell M. Unifying approaches from statistical genetics and phylogenetics for mapping phenotypes in structured populations. bioRxiv. 2024;.

15. Aw AJ, McRae J, Rahmani E, Song YS. Highly parameterized polygenic scores tend to overfit to population stratification via random effects; 2024. https://www.biorxiv.org/content/10.1101/2024.01.27.577589v1.

16. Veller C, Coop G. Interpreting population and family-based genome-wide association studies in the presence of confounding; 2023. https://www.biorxiv.org/content/10.1101/2023.02.26.530052v1.

17. Mostafavi H, Harpak A, Agarwal I, Conley D, Pritchard JK, Przeworski M. Variable prediction accuracy of polygenic scores within an ancestry group. Elife. 2020;9.

18. Howe LJ, Nivard MG, Morris TT, Hansen AF, Rasheed H, Cho Y, et al. Within-sibship genome-wide association analyses decrease bias in estimates of direct genetic effects. Nat Genet. 2022;54(5):581–592.

19. Tan T, Jayashankar H, Guan J, Nehzati SM, Mir M, Bennett M, et al. Family-GWAS reveals effects of environment and mating on genetic associations. medRxiv. 2025;.

20. Smith SP, Smith OS, Mostafavi H, Peng D, Berg JJ, Edge MD, et al. A litmus test for confounding in polygenic scores. bioRxivorg. 2025;.

21. Bycroft C, Freeman C, Petkova D, Band G, Elliott LT, Sharp K, et al. The UK Biobank resource with deep phenotyping and genomic data. Nature. 2018;562(7726):203–209.

22. Chen M, Sidore C, Akiyama M, Ishigaki K, Kamatani Y, Schlessinger D, et al. Evidence of polygenic adaptation in Sardinia at height-associated loci ascertained from the Biobank Japan. Am J Hum Genet. 2020;107(1):60–71.

23. Ding Y, Hou K, Xu Z, Pimplaskar A, Petter E, Boulier K, et al. Polygenic scoring accuracy varies across the genetic ancestry continuum. Nature. 2023; p. 1–8. 10.1038/s41586-023-06079-4.

24. Cook JP, Mahajan A, Morris AP. Fine-scale population structure in the UK Biobank: implications for genome-wide association studies. Human Molecular Genetics. 2020;29(16):2803–2811. 10.1093/hmg/ddaa157.

25. Bulik-Sullivan BK, Loh PR, Finucane HK, Ripke S, Yang J, Schizophrenia Working Group of the Psychiatric Genomics Consortium, et al. LD Score regression distinguishes confounding from polygenicity in genome-wide association studies. Nat Genet. 2015;47(3):291–295.

26. Devlin B, Roeder K. Genomic Control for Association Studies. Biometrics. 1999;55(4):997–1004. 10.1111/j.0006-341X.1999.00997.x.

27. Koenig Z, Yohannes MT, Nkambule LL, Zhao X, Goodrich JK, Kim HA, et al. A harmonized public resource of deeply sequenced diverse human genomes. Genome Res. 2024;34(5):796–809.

28. Bycroft C, Freeman C, Petkova D, Band G, Elliott LT, Sharp K, et al. The UK Biobank resource with deep phenotyping and genomic data. Nature. 2018;562(7726):203–209. 10.1038/s41586-018-0579-z.

29. Turchin MC, Chiang CW, Palmer CD, Sankararaman S, Reich D, Hirschhorn JN. Evidence of widespread selection on standing variation in Europe at height-associated SNPs. Nature Genetics. 2012;44(9):1015–1019. 10.1038/ng.2368.

30. Berg JJ, Coop G. A Population Genetic Signal of Polygenic Adaptation. PLoS Genetics. 2014;10(8):e1004412. 10.1371/journal.pgen.1004412.

31. Josephs EB, Berg JJ, Ross-Ibarra J, Coop G. Detecting Adaptive Differentiation in Structured Populations with Genomic Data and Common Gardens. Genetics. 2019;211(3):989–1004. 10.1534/genetics.118.301786.

32. Abdellaoui A, Hugh-Jones D, Yengo L, Kemper KE, Nivard MG, Veul L, et al. Genetic cor-relates of social stratification in Great Britain. Nature Human Behaviour. 2019;3(12):1332–1342. 10.1038/s41562-019-0757-5.

33. Akbari A, Barton AR, Gazal S, Li Z, Kariminejad M, Perry A, et al. Pervasive findings of directional selection realize the promise of ancient DNA to elucidate human adaptation. bioRxiv. 2024;.

34. Knowler WC, Williams RC, Pettitt DJ, Steinberg AG. Gm3;5,13,14 and type 2 diabetes mellitus: an association in American Indians with genetic admixture. American Journal of Human Genetics. 1988;43(4):520–526.

35. Lander ES, Schork NJ. Genetic dissection of complex traits. Science (New York, NY). 1994;265(5181):2037–2048. 10.1126/science.8091226.

36. Browning SR, Browning BL. Population Structure Can Inflate SNP-Based Heritability Estimates. The American Journal of Human Genetics. 2011;89(1):191–193. 10.1016/j.ajhg.2011.05.025.

37. Bulik-Sullivan B, Finucane HK, Anttila V, Gusev A, Day FR, Loh PR, et al. An atlas of genetic correlations across human diseases and traits. Nature Genetics. 2015;47(11):1236–1241. 10.1038/ng.3406.

38. Lewis ACF, Molina SJ, Appelbaum PS, Dauda B, Di Rienzo A, Fuentes A, et al. Getting genetic ancestry right for science and society. Science. 2022;376(6590):250–252. 10.1126/science.abm7530.

39. Steiner MC, Rice DP, Biddanda A, Ianni-Ravn MK, Porras C, Novembre J. Study design and the sampling of deleterious rare variants in biobank-scale datasets. bioRxivorg. 2025;.

40. Patterson N, Price AL, Reich D. Population Structure and Eigenanalysis. PLOS Genetics. 2006;2(12):e190. 10.1371/journal.pgen.0020190.

41. Bryc K, Bryc W, Silverstein JW. Separation of the largest eigenvalues in eigenanalysis of genotype data from discrete subpopulations. Theoretical Population Biology. 2013;89:34–43. 10.1016/j.tpb.2013.08.004.

42. Zoledziewska M, Sidore C, Chiang CWK, Sanna S, Mulas A, Steri M, et al. Height-reducing variants and selection for short stature in Sardinia. Nature Genetics. 2015;47(11):1352–1356. 10.1038/ng.3403.

43. Chen M, Sidore C, Akiyama M, Ishigaki K, Kamatani Y, Schlessinger D, et al. Evidence of Polygenic Adaptation in Sardinia at Height-Associated Loci Ascertained from the Biobank Japan. The American Journal of Human Genetics. 2020;107(1):60–71. 10.1016/j.ajhg.2020.05.014.

44. The Pan-UKB Team. Pan-UK Biobank: Study Design; 2020. https://pan.ukbb.broadinstitute.org/docs/study-design.

45. Shemirani R, Belbin GM, Cullina S, Caggiano C, Gignoux C, Zaitlen N, et al. SPC: a SPectral Component approach leveraging Identity-by-Descent graphs to address recent population structure in genomic analysis; 2025. https://www.medrxiv.org/content/10.1101/2025.06.04.25328990v2.

46. Zaidi AA, Mathieson I. Demographic history impacts stratification in polygenic scores. Genetics; 2020. http://biorxiv.org/lookup/doi/10.1101/2020.07.20.212530.

47. Hu S, Ferreira LAF, Shi S, Hellenthal G, Marchini J, Lawson DJ, et al. Fine-scale population structure and widespread conservation of genetic effect sizes between human groups across traits. Nat Genet. 2025;57(2):379–389.

48. Fan C, Mancuso N, Chiang CWK. A genealogical estimate of genetic relationships. bioRxiv. 2021; p. 2021.08.18.456747.

49. Johnstone IM, Paul D. PCA in High Dimensions: An Orientation. Proceedings of the IEEE. 2018;106(8):1277–1292. 10.1109/JPROC.2018.2846730.

50. Lango Allen H, Estrada K, Lettre G, Berndt SI, Weedon MN, Rivadeneira F, et al. Hundreds of variants clustered in genomic loci and biological pathways affect human height. Nature. 2010;467(7317):832– 838. 10.1038/nature09410.

51. Wood AR, Esko T, Yang J, Vedantam S, Pers TH, Gustafsson S, et al. Defining the role of common variation in the genomic and biological architecture of adult human height. Nature Genetics. 2014;46(11):1173–1186. 10.1038/ng.3097.

52. Howe LJ, Nivard MG, Morris TT, Hansen AF, Rasheed H, Cho Y, et al. Within-sibship genome-wide association analyses decrease bias in estimates of direct genetic effects. Nature Genetics. 2022;54(5):581–592. 10.1038/s41588-022-01062-7.

53. Smith WCS, Tunstall-Pedoe H, Crombie IK, Tavendale R. Concomitants of Excess Coronary Deaths — Major Risk Factor and Lifestyle Findings from 10,359 Men and Women in the Scottish Heart Health Study. Scottish Medical Journal. 1989;34(6):550–555. 10.1177/003693308903400603.

54. Privé F, Arbel J, Vilhjálmsson BJ. LDpred2: better, faster, stronger. Bioinformatics. 2021;36(22-23):5424–5431.

55. Ge T, Chen CY, Ni Y, Feng YCA, Smoller JW. Polygenic prediction via Bayesian regression and continuous shrinkage priors. Nat Commun. 2019;10(1):1776.

56. Spence JP, Sinnott-Armstrong N, Assimes TL, Pritchard JK. A flexible modeling and inference frame-work for estimating variant effect sizes from GWAS summary statistics; 2022. https://www.biorxiv.org/content/10.1101/2022.04.18.488696v1.

57. Song S, Benonisdottir S, Liu JS, Kong A. Participation bias in the estimation of heritability and genetic correlation. Proceedings of the National Academy of Sciences. 2025;122(25):e2425530122. 10.1073/pnas.2425530122.

58. Robinson MR, Kleinman A, Graff M, Vinkhuyzen AAE, Couper D, Miller MB, et al. Genetic evidence of assortative mating in humans. Nat Hum Behav. 2017;1(1):0016.

59. Kong A, Thorleifsson G, Frigge ML, Vilhjalmsson BJ, Young AI, Thorgeirsson TE, et al. The nature of nurture: Effects of parental genotypes. Science. 2018;359(6374):424–428. 10.1126/science.aan6877.

60. Belsky DW, Harden KP. Phenotypic Annotation: Using Polygenic Scores to Translate Discoveries From Genome-Wide Association Studies From the Top Down. Current Directions in Psychological Science. 2019;28(1):82–90. 10.1177/0963721418807729.

61. Chang CC, Chow CC, Tellier LC, Vattikuti S, Purcell SM, Lee JJ. Second-generation PLINK: rising to the challenge of larger and richer datasets. Gigascience. 2015;4(1):7.

62. Maier R, Flegontov P, Flegontova O, Işildak U, Changmai P, Reich D. On the limits of fitting complex models of population history to f-statistics. Elife. 2023;12.

63. Mbatchou J, Barnard L, Backman J, Marcketta A, Kosmicki JA, Ziyatdinov A, et al. Computationally efficient whole-genome regression for quantitative and binary traits. Nat Genet. 2021;53(7):1097–1103.

64. McVean G. A genealogical interpretation of principal components analysis. PLoS Genet. 2009;5(10):e1000686.

65. Engelhardt BE, Stephens M. Analysis of Population Structure: A Unifying Framework and Novel Methods Based on Sparse Factor Analysis. PLOS Genetics. 2010;6(9):e1001117. 10.1371/journal.pgen.1001117.

